# Generalized hippocampal replay exhibits memory recall-like dynamics

**DOI:** 10.64898/2026.11.12.766222

**Authors:** George Cai, Brian Lustig, Albert K Lee, Marcelo G Mattar, Samuel J Gershman

## Abstract

Hippocampal replay has been intensely studied for theorized roles in memory consolidation, credit assignment in reinforcement learning, and online planning. Much work in the field assumes that replayed trajectories traverse states smoothly and sequentially, in either forward or reverse order of experience. In a synthetic setting, we demonstrate that incorrect transition priors lead to systematic decoding error and show how model comparison addresses this issue. We compare a diverse set of transition models, including recall-like transition dynamics, and show how they can capture qualitatively different replay structures. Applying model comparison to three neural recording datasets from adult male rats, we find that the recall transition dynamics offer a statistically superior description of spiking during population burst events compared to a commonly used heuristic model of smooth trajectories. We propose a generalized conception of replay events through this theoretical connection to human free recall dynamics. Furthermore, our theory suggests that generalized replays may be a neural instantiation of online planning through generalized rollouts. We find preliminary evidence that generalized replays are more prevalent in settings where planning is beneficial. Overall, the proposed theory has implications for decoding methodology and the normative interpretation of hippocampal replay.

## 1. Introduction

The hippocampus has notable functions in forming episodic memories (Scoville and Milner, 1957; Morris et al., 1982) and representing a cognitive map (O’Keefe and Nadel, 1978). A large class of hippocampal neurons are functionally termed place cells because they track an animal’s position as it traverses through space during behavior. Nevertheless, structured, spontaneous activity has been observed in place cell ensembles while the animal is outside the associated place fields. After experience, the spontaneous activity showed sequential structure that was correlated to spiking during behavior, so the events were termed “replay” (Wilson and McNaughton, 1994; Skaggs and McNaughton, 1996; Lee and Wilson, 2002). The activity was assumed to be a literal replay of experience, so continuous spatial trajectories during experience implied that replayed trajectories should also be spatially continuous. This memory-like interpretation was consistent with the complementary learning systems theory, where the fast learning hippocampus would gradually “teach” cortex by replaying experiences (McClelland et al., 1995; Carr et al., 2011).

Subsequent work using Bayesian decoders on place cell population recordings showed that replayed sequences occurred in both forward and reverse temporal order relative to the real experience, leading to functional interpretations of credit assignment for reinforcement learning and planning future trajectories (Foster and Wilson, 2006; Diba and Buzsáki, 2007; Karlsson and Frank, 2009; Pfeiffer and Foster, 2013; Mattar and Daw, 2018). However, recent studies of spatial replays in rodents have revealed that the “replayed” sequences do not simply recapitulate experience (Davidson et al., 2009; Gupta et al., 2010; Pfeiffer and Foster, 2013), nor do the encoded trajectories smoothly traverse space (Pfeiffer and Foster, 2015; Berners-Lee et al., 2022). Instead, the decoded states within events exhibit jumps to discrete locations interleaved with hovering in place. These observations indicate that we should consider a broader concept of replay (Figure 1**A**). We will use the term “generalized replay” to refer to this broader class of sequentially structured neural activity for continuity with past literature, with the important distinction that the activity sequence during experience is not being replayed in a literal sense. Many traditional replay analyses have relied on memoryless, uniform-prior Bayesian decoding with scoring criteria that assume replays encode smooth, directed trajectories through space without jumps or reversals in direction. This style of analysis is biased against identifying generalized replay events. Our goal is to develop an analysis framework that can accurately identify generalized replay events, and to consider the theoretical implications of such events.

**Figure 1:**
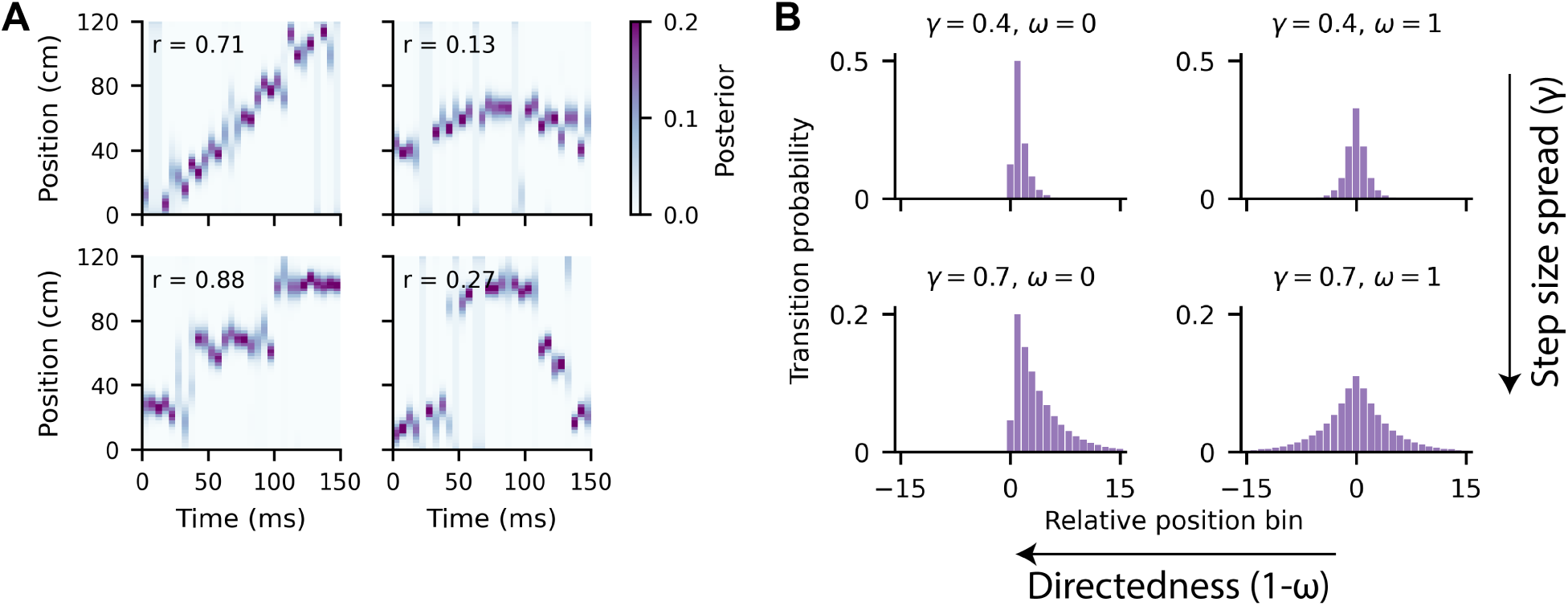
Exemplifying a range of different replay dynamic structures. **A** We simulated spikes and Bayesian decoding posterior heatmaps from 30 idealized place cells to illustrate assumptions about smoothness and directedness of replayed trajectories. Spatial correlation between posterior mean and time are annotated in each panel. Only the top left event is traditionally considered a replay event by satisfying both high spatial correlation and small step sizes. **B** The TCM-SR transition kernel can be parametrically varied in its step size spread and directedness. The parameter *ω ∈* [0, 1] modulates the directedness or asymmetry, and the discount factor *γ ∈* [0, 1) modulates the step size spread.

Recent studies have utilized state space models with priors on state transition dynamics to decode at finer temporal resolution and exclude fewer events. Assuming hidden state transitions are Markovian, transition kernels have two major axes of variation: step size spread and skewness. Step size spread controls the smoothness of decoded trajectories, with narrower spreads producing more continuous trajectories. Earlier transition kernels and analyses often explicitly or implicitly induce smoothness. Skewness affects how directed a trajectory is, and strongly skewed kernels produce trajectories that progress in a preferred direction. Two recent studies have used stationary, fragmented, and continuous modes, and diffusive trajectories with momentum (Denovellis et al., 2021; Krause and Drugowitsch, 2022). The continuous and momentum kernels are symmetric, that is, unskewed. Moreover, the relatively narrow distributions of continuous and diffusion with momentum kernels are intuitively inconsistent with the jumps in the empirically observed jump-and-hover structure. A mismatch between the decoder and data could introduce systematic decoding errors, although this has not been characterized. Lastly, from a theoretical standpoint, the previously chosen transition priors are heuristic and lack normative interpretations.

To address these gaps, this study explores a novel method for replay decoding that accommodates variation in both step-size spread and skewness. We first use simulated data under different ground-truth transition models to illustrate how mismatched priors can distort decoded trajectories. Then, we introduce a straightforward cross-validated model comparison framework to evaluate transition priors and select the kernel that minimizes decoding errors. Finally, we use our approach on three empirical datasets of awake hippocampal replay. Because no prior transition kernels are clearly consistent with the empirically observed jump-and-hover dynamics, we additionally propose a theoretically motivated transition kernel with independent parameters for step size spread and skewness (Figure 1**B**). Beyond parametric flexibility, the model connects replay dynamics to human memory of temporal sequences and reinforcement learning.

To motivate our theoretically derived kernel, we turn to human free recall of lists of memoranda, a classic episodic memory task that is hippocampally dependent (Manns et al., 2003; Vargha-Khadem et al., 1997). During free recall, the transition probabilities between retrieved items have a characteristic contiguous, forward-biased distribution (Kahana, 1996). These results were explained by the Temporal Context Model (TCM) and related models (Howard and Kahana, 2002; Sederberg et al., 2008; Polyn et al., 2009; Lohnas et al., 2015).

The critical mechanism of TCM is a drifting context vector that integrates presented stimuli. Bidirectional associations between stimuli and context formed during the encoding stage enable context-similarity based retrieval of memories. The model produces the contiguity effect, where recalling a stimulus makes stimuli that occurred nearby in time more likely to be retrieved next, and forward asymmetry, where subsequent stimuli in the original presentation order are more likely to be recalled than preceding stimuli (Howard and Kahana, 2002). Temporal context effects on recall predicted by this model have been observed in human hippocampal and amygdala single unit, fMRI and electroencephalography recordings (Folkerts et al., 2018; Bornstein and Norman, 2017; Rait et al., 2025; Manning et al., 2011; Gershman et al., 2013). Temporal representations at a variety of time scales have also been observed in rodent and non-human primate hippocampus and entorhinal cortex (Salz et al., 2016; MacDonald et al., 2011; Manns et al., 2007; Rueckemann et al., 2025; Tsao et al., 2018; Kanter et al., 2025).

Subsequent work normatively grounded TCM in a sequence prediction framework through a connection to the successor representation (SR). Given a state transition matrix, the SR matrix encodes the expected discounted future state visits (Dayan, 1993). In other words, it is a predictive map of future state occupancy that enables behavioral flexibility under changing rewards or goals in the same environment. Hippocampal CA1 place cells are thought to encode the SR (Stachenfeld et al., 2017), and behavioral evidence also supports the use of the SR in human reinforcement learning and decision making (Momennejad et al., 2017; Kahn et al., 2025). Under specific assumptions, there is a mathematical equivalence between the SR matrix and TCM’s context-item association matrix that is used to sample the next recalled item during retrieval (Gershman et al., 2012). Conceptually, the drifting temporal context can be seen as an eligibility trace for assigning credit to past states for being predictive of the current state; these predictive relationships are cached in the SR or context-item association matrix. We provide a detailed description of the theory in Section 4.1. Under this framework, free recall can be interpreted as iteratively sampling from the successor states of the current context, with higher probability for future states with higher expected occupancy. The context vector is simply a drifting mixture of states, so the retrieval distribution is a mixture of the successors of different states. The recalled samples can support online evaluation and decision-making in sequential decision problems (Zhou et al., 2025).

The explicit hypothesized connections between TCM-SR and replay are the following: (1) discrete hover locations in jump-and-hover replay dynamics correspond to discrete recalled memoranda; (2) jumping to new hover locations corresponds to sampling from a mixture of SRs; (3) jumps can transition forward or backward relative to the sequence of experienced states in a single replay event; and (4) jump probability follows the characteristic forward-biased contiguous distribution of free recall. Under this model of replay dynamics, events that were previously classified as non-replays or fragmented replays can be parsimoniously explained.

To evaluate transition models from prior work and our novel hypothesis, we reanalyzed rat hippocampal neural recordings to characterize the replayed state transitions during population burst events (Lustig et al., 2025; Grosmark and Buzsáki, 2016; Pfeiffer and Foster, 2015). Using both hidden Markov models (HMMs) and memoryless Bayesian decoding for model comparison, we find that decoded replay transitions are more consistent with TCM-SR in a memory- and planning-demanding task. We show how relaxing or modifying prior analysis assumptions about replay dynamics leads to quantitatively superior statistical descriptions of spiking during putative replay events, enabling us to test the TCM-SR theory’s predictions. The improved fit of our theoretical model of transition dynamics in a task with memory and planning demands justifies a more general conceptualization of replay structure and motivates the development of improved replay decoding methodology.

## 2. Results

Because there is no ground truth for replayed states in data, we first take a simulation approach to characterize how transition priors influence decoding error from generated spikes. We demonstrate that the HMM held-out likelihood is a good proxy for decoding error. Then, we use a model comparison approach to assess different transition priors (kernels) on neural data by held-out likelihood. Finally, we provide illustrative examples of how the resulting decoded trajectories differ between kernels.

Next, we provide an overview of ten alternative models of transition dynamics described in Table 2. Stationary, continuous, and fragmented kernels have been adopted in the hierarchical HMM of (Denovellis et al., 2021). Diffusion with and without momentum has been characterized in 2D arenas (Stella et al., 2019; Krause and Drugowitsch, 2022). These kernels are depicted in Figure 2**A** (first row). We include an additional control kernel termed segment uniform which allows uniform transition probability within a directional map. In this study, we introduce kernels derived from TCM-SR for comparison (Figure 2**A**, second row). Movement refers to the kernel that describes the empirical spatial bin transitions at 100 ms, approximately the theta oscillation period. SR and TCM-SR kernels use the successor representation of this movement transition matrix. In TCM-SR, the parameter *ω* ∊ [0, 1] modulates the directedness or asymmetry, with *ω* = 1 being a fully symmetric kernel and *ω* = 0 being a maximally forward-biased kernel, and the discount factor *γ* ∊ [0, 1) modulates the step size spread, with *γ* = 0 being a movement-like, one-step transition and *γ* > 0 corresponding to an SR-like, multi-step prediction of future states with an exponential discount factor. Lastly, we include reversed SR and symmetrized TCM-SR kernels as additional controls, although the TCM-SR kernel can express the SR (*ω* = 0*, γ >* 0), symmetrized TCM-SR (*ω* = 1*, γ >* 0), and movement kernels (*ω* = 0*, γ* = 0) as special cases of parameter choice. Thus, finding fitted values 0 < *γ* < 1 and 0 < *ω* < 1 is evidence of TCM-SR-like dynamics rather than a simpler model in this family.

**Figure 2:**
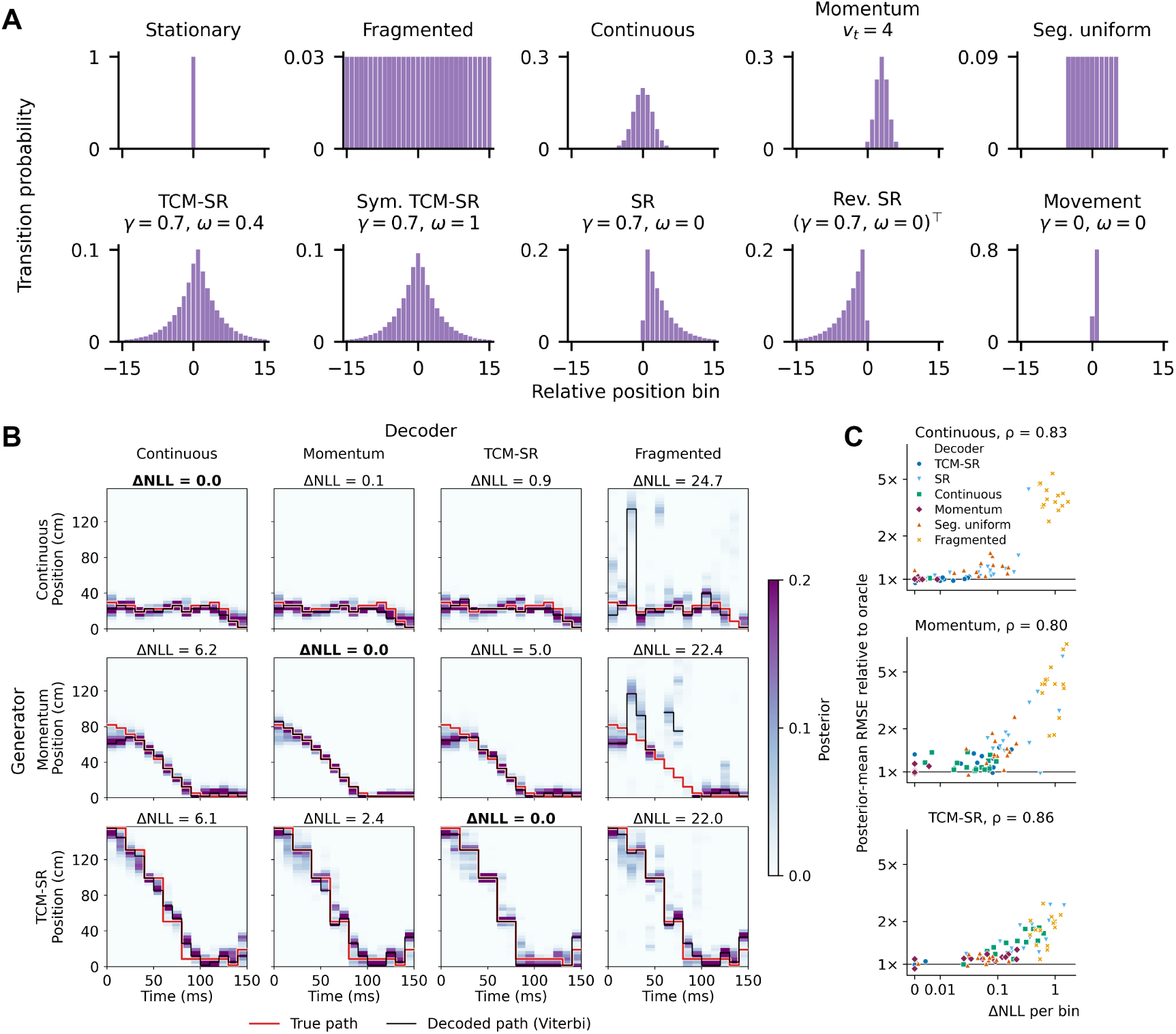
Mismatched transition priors result in decoding errors. Using place fields derived from experimental recordings in (Lustig et al., 2025), we simulate time bin-matched spiking data with ground truth hidden state trajectories according to different dynamics. **A** Schematized transition probability distribution to relative positions for 10 kernels. The mean offset of momentum is time varying. The second row’s kernels are all expressible with the TCM-SR kernel or its transpose using different parameters. **B** Example trajectories generated by continuous, momentum, and TCM-SR dynamics (rows) are shown in red traces. Each column represents a fitted decoder on a training split of the data, with heatmaps showing marginal posterior and black traces showing the most likely sequence of states (Viterbi path). The fragmented decoder’s Viterbi path is discontinuous because the most probable bin was on another map. ΔNLL is the difference in negative log-likelihood under the decoder model relative to the oracle, and closer to 0 is better fit. **C** Posterior-mean RMSE relative to the generative model with true parameters plotted against ΔNLL per bin for different fitted models. Each point represents a simulated session using place fields and generating parameters fitted from a real session’s data. *ρ* denotes Spearman correlation.

### 2.1. Decoding errors are introduced by mismatched transition priors

To systematically evaluate the cost of a mismatched transition prior, we sample spike trains with ground truth hidden state trajectories and compute the decoding error from fitted HMMs with different transition priors. The observation model for spike counts was conditionally independent Poisson spiking with rates determined by place fields (see Eq. 17) from the experimental data of (Lustig et al., 2025), and the number of time bins simulated was matched to the duration of experimentally identified events. Thus, the only differences between the generative HMM and decoder come from the transition model class and parameters. The generative model’s transition parameters were uniformly sampled from the parameter space. Then, we fit parameters in decoding HMMs using a generalized Expectation Maximization (EM) algorithm to maximize the likeli-hood of the simulated binned spike counts. Finally, we decode held-out spikes using the forward-backward algorithm and compute the error.

We illustrate the errors introduced by mismatched transition priors with example trajectories and decoding heatmaps shown in Figure 2**B** for a subset of kernels. We focus on this subset because the continuous and momentum kernels are emphasized in prior work and encode the assumption that replay of spatial trajectories is veridical. By contrast, TCM-SR produces non-veridical, jumpy trajectories. The three rows depict different generating models. The fragmented transition model in the fourth column is a special case of the HMM framework that is equivalent to a memoryless Bayesian decoder with uniform prior. Evidently, different transition priors influence the smoothed posterior and Viterbi (maximum likelihood) trajectory. In the first column, the continuous kernel artificially smooths the trajectory generated by the TCM-SR kernel, and in the second row, TCM-SR and continuous kernels shift the initial position of the trajectory generated by the momentum kernel.

Aggregating across all events and sessions, we find that kernel mismatch can increase decoding error. We compute the root mean-squared error (RMSE) of the mean of the decoder’s posterior distribution in each time bin and compare to the original generating model’s RMSE in Table 1. If the generating transition model is TCM-SR or momentum, then kernel mismatches result in worse decoding performance even after parameter fitting. The data generated by the continuous kernel are accurately decoded by fitted momentum and TCM-SR kernels. This is sensible because the continuous kernel is a special case of momentum and is well-approximated by TCM-SR if the spread is narrow. However, we cannot measure decoding error in real neural data since there are no ground truth latent states, so we use the decoder marginal log likelihood as a proxy measure (Figure 2**C**). Formally, log likelihood quantifies how well a model explains the observed data, and we find robust correlations between the difference in log likelihood from the best fitting model (which matches the oracle in our synthetic setting) and the decoding error. Thus, we can evaluate transition models via the likelihood on held-out data when analyzing neural recordings.

**Table 1:** Decoding error under different transition kernels across generator parameter space. In 14 sessions with different place fields, each generator was simulated at 100 random tuples uniformly sampled throughout the parameter space of each model. The stickiness *κ* is each session’s fitted value. Entries are the RMSE of decoder’s posterior-mean position on held-out events relative to that of the oracle noise floor (the generating kernel at its true parameters). We show median over all settings with interquartile range below, and 1.00 is oracle-level. Decoders were fitted on 80% of events and evaluated on held-out 20%.

|  | Decoder |  |  |  |  |  |  |
|---|---|---|---|---|---|---|---|
| Generator | Continuous | Momentum | TCM-SR | SR | Movement | Seg. uniform | Fragmented |
| Continuous | 1.00<br>(1.00–1.01) | 1.00<br>(1.00–1.01) | 1.02<br>(1.00–1.04) | 1.17<br>(1.07–1.34) | 1.17<br>(1.07–1.32) | 1.17<br>(1.09–1.29) | 3.73<br>(3.10–4.49) |
| Momentum | 1.04<br>(1.01–1.15) | 1.00<br>(1.00–1.02) | 1.05<br>(1.01–1.12) | 1.41<br>(1.15–1.91) | 1.45<br>(1.16–1.99) | 1.25<br>(1.13–1.41) | 3.37<br>(2.50–4.50) |
| TCM-SR | 1.14<br>(1.04–1.35) | 1.09<br>(1.03–1.18) | 1.00<br>(1.00–1.01) | 1.14<br>(1.03–1.43) | 1.29<br>(1.07–1.71) | 1.17<br>(1.08–1.33) | 2.69<br>(1.94–3.61) |
| SR | 1.26<br>(1.13–1.52) | 1.20<br>(1.10–1.35) | 1.01<br>(1.00–1.03) | 1.00<br>(0.99–1.01) | 1.15<br>(1.01–1.56) | 1.31<br>(1.18–1.50) | 3.53<br>(2.85–4.44) |
| Movement | 1.22<br>(1.13–1.31) | 1.15<br>(1.08–1.25) | 1.01<br>(1.00–1.02) | 1.00<br>(1.00–1.01) | 1.00<br>(1.00–1.01) | 1.53<br>(1.33–1.76) | 4.54<br>(3.79–5.66) |

### 2.2. TCM-SR transition dynamics best explain neural data in a memory- and planning-demanding task

To compare the dynamics models, we analyzed neural recordings from (Lustig et al., 2025; Grosmark and Buzsáki, 2016; Pfeiffer and Foster, 2015). Briefly, (Lustig et al., 2025) trained rats to perform a 3-arm alternation task on a W-maze to collect water rewards at the end of each arm (Figure S1**A**). Between each arm run, the subjects were required to run on a wheel for 8 seconds to open doors from a central hub to the arms. Subjects were trained to expert behavioral performance before neural recording. In (Grosmark and Buzsáki, 2016), rats were trained to collect water reward on an open-field cheeseboard maze. Then, they were introduced to novel linear tracks in unfamiliar rooms where water reward was collected by alternating between the ends of the track (Figure S1**B**). In (Pfeiffer and Foster, 2015), rats were trained to collect liquid food reward by alternating between ends of a 180 cm linear track. Neural data from awake, on-task epochs of all three datasets were analyzed.

The TCM-SR model family can encode asymmetric priors, which requires us to define the forward direction in a given replay event. We leverage the fact that many CA1 cells exhibit different place fields in each direction of travel in linear environments. Following (Lustig et al., 2025), we use the terms “outbound” and “inbound” to identify the physical direction of travel that a map corresponds to while “forward” and “reverse” describe the tendency of replay displacements relative to the active map. In the W-maze, there are three arms with two directional place maps each. A seventh map is included for the wheel, with states defined as 125 ms time bins running on the wheel. To determine which directional map was active during events, we compute posterior probabilities over spatial states that summed to 1 over all maps simultaneously (seven maps for (Lustig et al., 2025) and two maps for (Grosmark and Buzsáki, 2016; Pfeiffer and Foster, 2015)).

We evaluate transition models on three sets of event inclusion criteria to ensure our comparison results are robust. First, we identify “traditional replays” using a memory-less Bayesian decoder with uniform prior and thresholding based on weighted Spearman correlation and minimum duration within population burst events (PBEs). Second, we relax the requirement on spatial correlation and allow events with high posterior mass on the same map in consecutive time bins, which we term “same-map activation events” (see Methods 4.5). Third, we conduct model comparison on all PBEs with spikes represented in non-overlapping 3 ms time bins. We present the results from analyzing same-map activation events throughout the main figures unless otherwise indicated to reduce spatial criteria on event selection.

Applying cross-validated likelihood comparison to fitted HMMs (see Methods 4.7), we find the TCM-SR or symmetrized TCM-SR transition kernel is the best fit on 13/18 sessions across 7 rats in the W-maze data (Figure 3**B**). In two linear track datasets, momentum dynamics are the best fit on 2/3 sessions across 3 rats in (Grosmark and Buzsáki, 2016), and 7/9 sessions across 3 rats in (Pfeiffer and Foster, 2015) (Figure S2**A-B**). Nevertheless, TCM-SR family kernels fit the data better than the continuous kernel in 2/3 sessions from (Grosmark and Buzsáki, 2016) and 9/9 sessions from (Pfeiffer and Foster, 2015). We excluded 3 sessions due to insufficient (< 10) events (see Tables S1, S2, S3 for event counts). The fitted discount factor *γ* and asymmetry parameter *ω* values indicate that the TCM-SR kernel was distinct from the SR (*ω* = 0), symmetrized TCM-SR (*ω* = 1) and movement transitions (*ω* = *γ* = 0) in many sessions (Figure S3). Moreover, the wide range of fitted values indicates the flexibility of the TCM-SR parameterization improves the kernel’s ability to match the data generating process. The continuous kernel’s *σ*_arm_ parameter for the track arms was saturated to the cap in nearly all sessions, indicating the local continuity assumption of replay transitions is not statistically supported.

**Figure 3:**
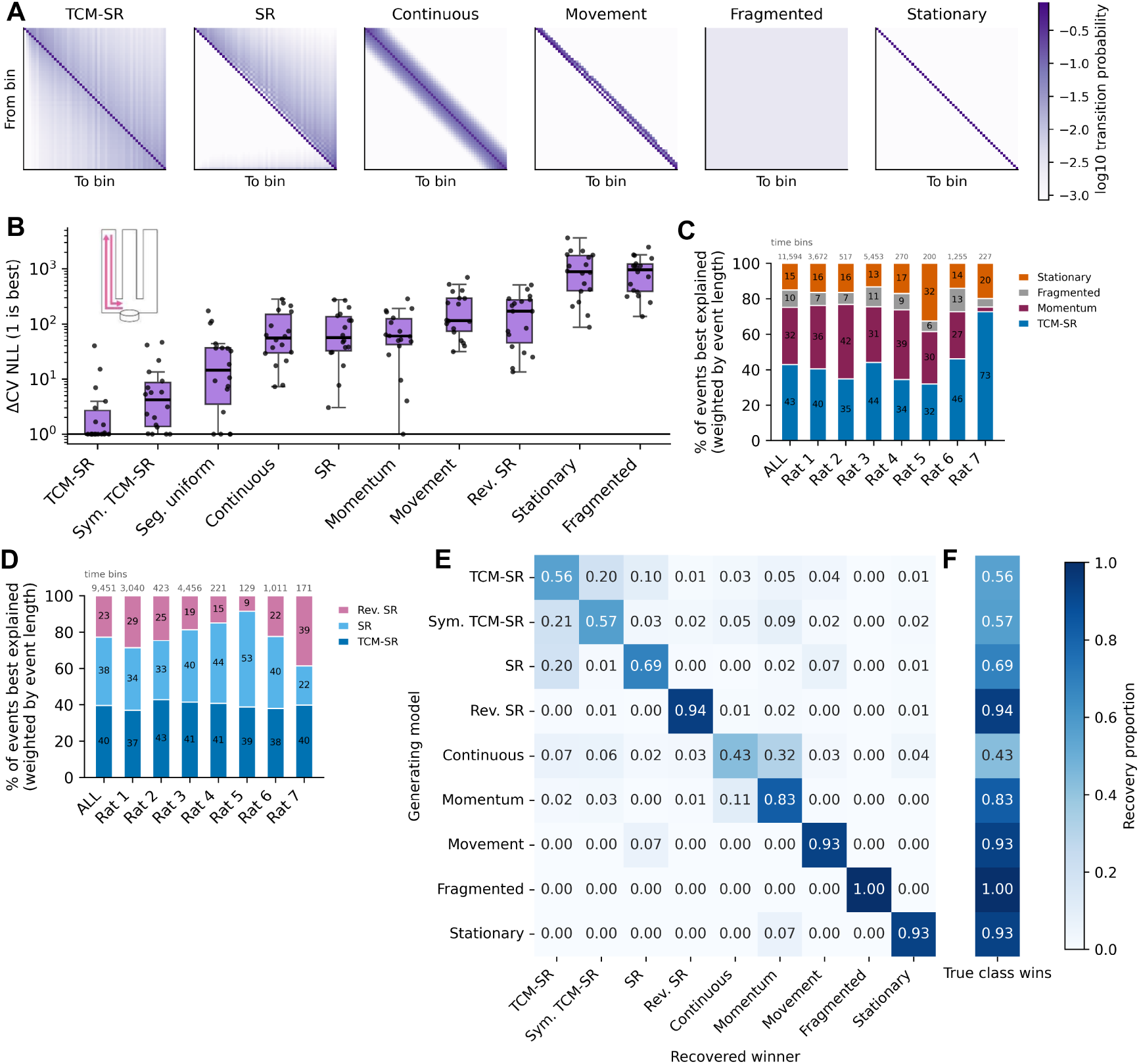
Replay dynamics are best explained by the TCM-SR transition model under model comparison in W-maze data. **A** Representative fitted transition kernels for a single directional map. **B** Difference in negative log-likelihood under 5-fold cross-validation at the event level; we add 1 for log scale display. Each point denotes a single session. **C** Fraction of events best explained by each of four transition kernels weighted by event duration. **D** Fraction of events best described by SR, reversed SR, and TCM-SR after removing events best described by stationary and fragmented kernels. **E** Recovery matrix of 5-fold cross-validation model comparison procedure using 1000 randomly sampled parameter tuples for each generating kernel. The amount of synthetic spiking data was matched to the experimental data on a time-bin basis. **F** Marginal recovery proportion for each generating model.

Because different events within a session may be consistent with different dynamics (Krause and Drugowitsch, 2022), we check that nearly half of all events were best explained by TCM-SR (Figure 3**C**). We omit special case and one sided kernels for a more direct comparison of TCM-SR and diffusion with momentum. We also find SR, TCM-SR, and reversed SR each best explain a comparable fraction of events when these three kernels are compared (Figure 3**D**).

Next, we check whether the lead of TCM-SR over continuous and momentum dynamics depended on event selection criteria. We find the result holds on traditional replay events in the W-maze dataset (Figure S4**A**; see Figure S2**C-D** for linear track). We use kernels fit on one session to compute likelihood on another session, and we find the model comparison result was robust to cross-session transfer within rats (Figure S4**B**). The improved fit of TCM-SR persisted when comparing dynamics on all PBEs without further filtering and decoding at 3 ms time bins (Figure S4**C**). In the linear track data, TCM-SR and momentum were comparably good descriptions of spiking during unfiltered PBE events at 3 ms time bins (Figure S2**E-F**). Thus, the model comparison was robustly in favor of TCM-SR over continuous and momentum models across a range of data filtering and binning settings for the W-maze data but not the linear track data.

We conducted a synthetic model recovery analysis to ensure the cross-validated model fitting and comparison procedure could identify data generated from each transition model. On representative sessions from 3 rats, we generated event duration-matched time bins of spiking data under each transition model with 1,000 uniformly randomly sampled parameters. We find robust recovery of the generating model (Figure 3**E-F**). The substantial off-diagonals for SR, Sym. TCM-SR, and TCM-SR generated data can be explained by the SR being a special case of TCM-SR with *ω* = 0 and Sym. TCM-SR being a special case with *ω* = 1. Likewise, the continuous kernel is a special case of the momentum kernel at the *a* → 0 limit, leading to mis-identification in that parameter regime. Recovery is generally consistent across parameter space, with the exception of strong self-transition probability (*κ* ∊ (0.79, 0.99)) resulting in non-identifiability because stickiness was a feature in all kernels except fragmented (Figure S5). In the linear track datasets, we find comparably robust model recovery in (Pfeiffer and Foster, 2015) but substantially weaker recovery in (Grosmark and Buzsáki, 2016) due to fewer units recorded and candidate events detected (Figure S6). Accordingly, the model comparison result for (Grosmark and Buzsáki, 2016) should be interpreted with caution. Overall, the recovery analysis shows the amount of data in the larger datasets is suitable for identifying different generative models using our model comparison method.

### 2.3. Replay jump distributions exhibit broad step size spread consistent with TCM-SR dynamics

Next, we ask which transition priors are consistent with state transitions observed in the data. We compute the histogram of decoded transitions in the maximum a posteriori (MAP) state with hovers omitted during same-map activation events. We refer to this histogram as the jump distribution. Using a memoryless Bayesian decoder with uniform prior, we find a clear contiguity effect and significant forward asymmetry in both traditional replay events and same-map activation events in the W-maze data (Figure 4**A-B**). To compare to transition priors, we sampled synthetic trajectories with matched sequence length to illustrate the fitted TCM-SR transition prior is consistent with the forward asymmetry and contiguity effects (Figure 4**C**). By contrast, the jumps generated by the momentum kernel exhibit locality and symmetry, which are qualitatively different from the jump distribution under the naive decoder (Figure 4**D**). In the linear track datasets, the naive decoder jump distribution (Figure S7**A-B**) also shows jump distribution spread that is qualitatively more consistent with TCM-SR or symmetrized TCM-SR than the momentum kernel (Figure S7**C-F**). However, the transition asymmetry is forward in (Grosmark and Buzsáki, 2016) and reversed in (Pfeiffer and Foster, 2015). We address possible explanations for the different asymmetry directions in the Discussion. Lastly, we compare the one-step transition probabilities from decoded neural data to the fitted kernels as a graphical posterior predictive check (Figure S8). We find that TCM-SR captures the bulk of the jump distribution decoded in the W-maze data and the linear track data from (Grosmark and Buzsáki, 2016), but all kernels deviate at intermediate jump distances in (Pfeiffer and Foster, 2015).

**Figure 4:**
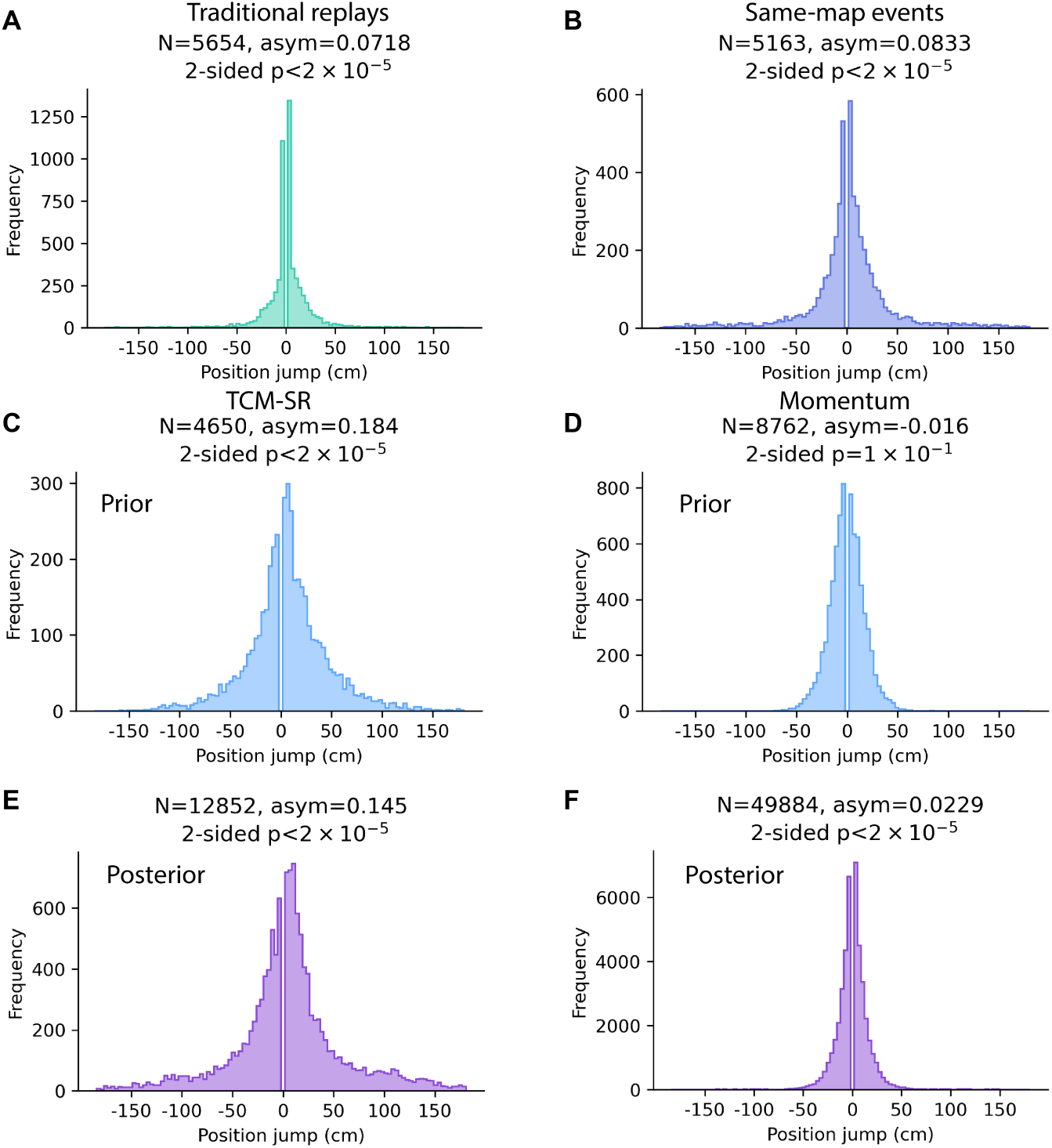
Jump distribution under memoryless Bayes, TCM-SR, and momentum decoders. Histogram of jumps in the decoded position between adjacent time bins during events. Transitions to the same bin are omitted as hovers. *N* denotes the number of jumps, “asym” is an asymmetry index with values in [ –1, 1], and 2-sided *p*-values were computed against a binomial null distribution for the asymmetry. **A** Jump distribution computed in replay events classified under traditional spatial correlation and sequence length criteria. We used 20 ms time bins stepped at 5 ms. **B** Jump distribution in replay events identified by a threshold on contiguous time bins being decoded to a single directional map. Event identification and decoding both used non-overlapping 10 ms time bins. **C-D** Generated jump distributions by sampling trajectories from the fitted TCM-SR and momentum kernels respectively. **E-F** Decoded jump distribution using fitted TCM-SR and momentum HMM from same-map activation events, respectively.

Finally, we ask how the posterior jump distribution under TCM-SR and momentum kernels compare. Using the Viterbi algorithm, we decode the sequence of most probable latent states in each PBE under fitted HMMs (Viterbi, 1967; Linderman et al., 2020). The jump distributions under fitted kernels are shown in Figure 4**E-F** (see Figure S9 for ratby-rat distributions). The TCM-SR decoded distribution exhibits heavier tails than the momentum kernel result, as expected from the predictive horizon of the SR. This matches the key characteristics of the naive memoryless Bayes decoded distribution (Figure 4**B**) and the prior. Thus, the memoryless Bayesian decoder with uniform prior, TCM-SR prior, and TCM-SR posterior all concur on the qualitative characteristics of a broad, forwardbiased jump distribution in the W-maze data.

### 2.4. Generalized replay avoids the need for heuristic selection criteria

How do the trajectories that are better fit by a TCM-SR transition prior differ from those better fit by a symmetric, local kernel? We show a representative selection of events with different log likelihood under the TCM-SR and momentum kernels in Figure 5. The final row uses the fragmented kernel, which is equivalent to naive memoryless Bayesian decoding with a uniform prior.

**Figure 5:**
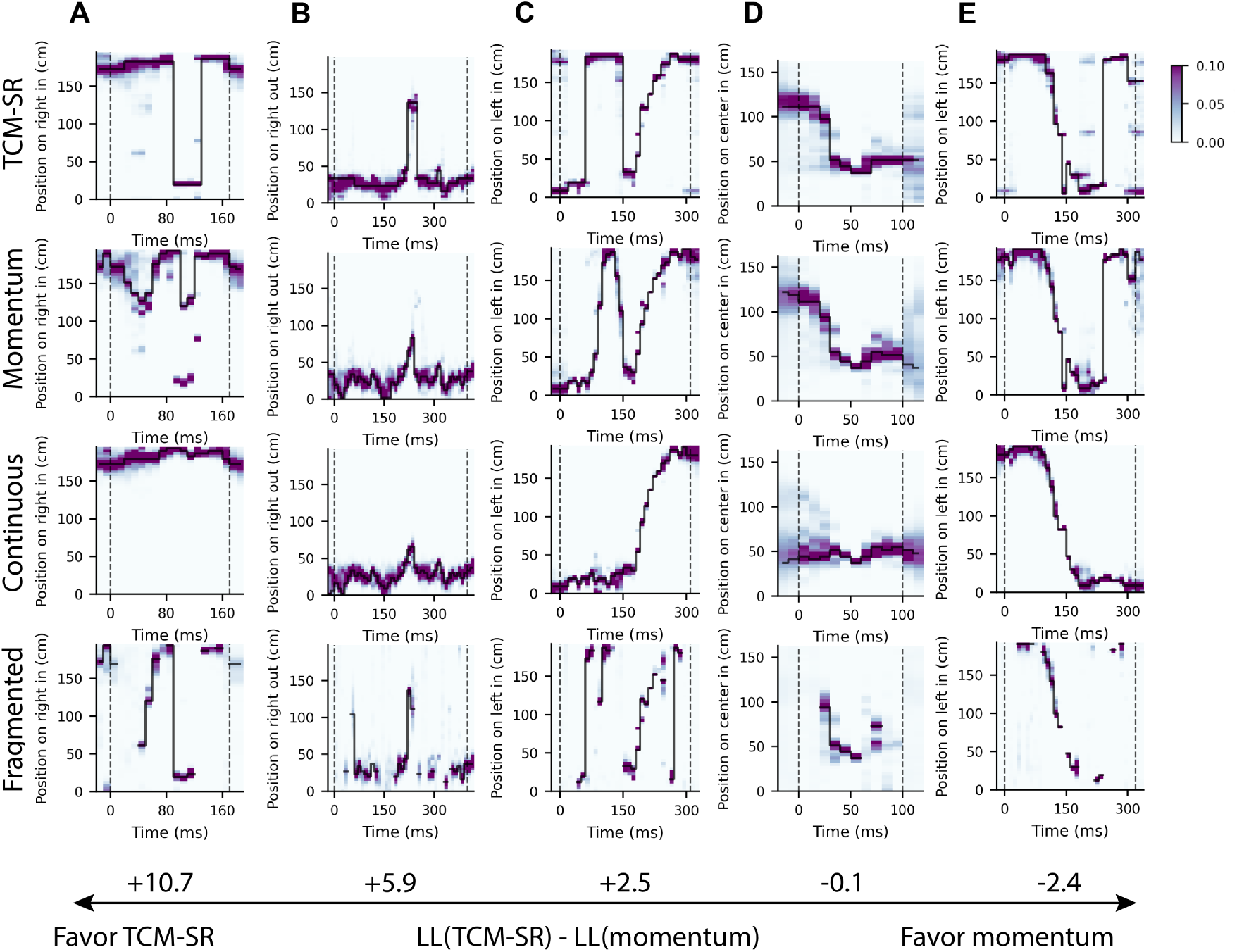
Generalized replay under different transition kernels. We show 5 examples of generalized replay events decoded by the TCM-SR, momentum, continuous, and fragmented kernels. The heat maps display the posterior probability over states in the active map of the trajectory event. The black trace indicates the most likely path decoded by the Viterbi algorithm. Positive values of ΔLL favor TCM-SR relative to the momentum kernel. We show the fragmented kernel decoder for comparison since it is equivalent to the naive memoryless Bayesian decoder with uniform prior.

Trajectories better explained by TCM-SR contain larger jumps (Figure 5**A-C**) than those better explained by the momentum kernel. Notably, the continuous kernel smooths over jump gaps that are approximately 40 ms in duration. Additionally, some generalized replays reverse directions within a single event despite activating the same directional map (Figure 5**C,E**). The event in column **D** has similar log likelihood under TCM-SR, momentum, and continuous kernels. The continuous kernel produces a distinct trajectory from the former two in the early part of the event, although the heatmap of the continuous kernel places some posterior mass on a trajectory similar to the other kernels. Finally, the event in column **E** is better explained by the momentum model. We see that the traversal across the map is relatively directed and smooth, and this is consistent with the posterior mass on this map according to the fragmented decoder as well. Since the TCM-SR kernel better explained spiking data under model comparison, events like the ones on the left half of the figure are more representative of PBEs than the typically assumed smoother, more monotonic trajectories on the right.

Therefore, we argue against the use of replay event scores that assume monotonic continuity such as spatial correlation and line-fitting. These results also imply that implicit smoothing in earlier analysis steps such as using overlapping decode time bins or jittering spike times may not be statistically justifiable. Instead, we advocate for using state space models with explicitly formulated dynamics assumptions in the form of transition priors. Then, we can directly adjudicate between analysis priors using measures such as cross-validated log likelihood. Using this framework in our study, we demonstrate that a generalized notion of replay is a superior description of a large amount of replay data during a memory- and planning-demanding task.

## 3. Discussion

Our study challenges common assumptions about replay dynamics and presents a computationally motivated transition kernel for accurately decoding generalized replay. First, we show that when decoder priors are mismatched with respect to the true generative process, events are either distorted or discarded for violating these assumptions. To address this issue, we encode transition priors in HMMs and show that the log likelihood of observed spiking data is a good proxy for accurate decoding in synthetic simulations. Second, we apply a cross-validated likelihood comparison procedure to evaluate different transition models on neural data. Beyond previously implemented kernels, we introduce the TCM-SR transition prior for replay dynamics that can independently modulate the jump spread with discount factor *γ* and directional preference with asymmetry factor *ω*. We find that TCM-SR dynamics are a better statistical description of the spiking data than the common heuristic of continuity on all datasets, supporting its use for more accurate state space decoding. TCM-SR is also a better description than momentum for events during a task involving memory and planning. Overall, our analyses lead us to propose a generalized notion of replays that may jump and reverse directions that departs from previous assumptions of veridical replay.

Our work builds on recent efforts in state space decoding to make transition model assumptions explicit, improve temporal resolution of decoding, and avoid rejecting events based on heuristic criteria. (Denovellis et al., 2021) developed a hierarchical HMM that allows switching between dynamic modes within events. However, the parameters in the continuous transition kernel are not fitted to data and induce distortions in the decoded trajectories if the parameters or transition structure are mis-specified. Our work addresses these concerns by fitting parameters during model comparison and recovery analysis. (Krause and Drugowitsch, 2022) modeled 2D latent dynamics underlying awake sharp wave ripple activity and showed a large majority of awake events in (Pfeiffer and Foster, 2013) have spatial trajectories with momentum. The result contrasts with asleep replays in 2D environments following Brownian motion dynamics (Stella et al., 2019). In our study, we found that TCM-SR was a better description than diffusion with momentum for replays in the W-maze but not the linear track datasets. Two alternative hypotheses about the source of discrepancy are differences in track geometry (rats may need to slow down at the turning points on the W-maze) or task demands (arm alternation requires memory). We found that the subset of generalized replay events on the center arm of the W-maze still favor TCM-SR dynamics over momentum (Figure S4**D**), implying track geometry is not the explanation. Thus, we hypothesize that tasks with memory demands induce TCM-SR replay dynamics, which can be tested in future studies that analyze datasets from different tasks.

Leveraging structure in state space decoding methods enables analyzing replay events at finer temporal resolution (3 ms vs. 20 ms time bins). Small time bins are especially relevant for analyses involving slow-gamma oscillations (25–50 Hz) (Pfeiffer and Foster, 2015). Decoded trajectories may also have improved spatial resolution by avoiding averaging over hover states that occur nearby in time. However, these advantages are contingent on the transition prior matching the generating process. Parameter fitting improves this match, and the flexible parameterization of TCM-SR can accommodate different degrees of step size spread and directedness, including a close approximation to the local continuous kernel. This flexibility is important given the wide range of fitted parameters observed across sessions in the neural data. Thus, we implemented the TCM-SR kernel and parameter fitting functionality in the hierarchical HMM package developed by (Denovellis et al., 2021) as a replacement of the heuristic local continuity kernel. Future work may explore increasing transition model flexibility further by extending to a second order Markov approximation of TCM-SR or a combination with momentum.

TCM-SR dynamics have computational implications for planning and decision making through generalized replays. Prior work has theorized that forward replay is involved in prospective planning (Pfeiffer and Foster, 2013) while reverse replay can mediate credit assignment after rewards or during rest (Foster and Wilson, 2006; Diba and Buzsáki, 2007; Ambrose et al., 2016; Mattar and Daw, 2018). These theories implicitly view replay as directed, smooth rollouts because that is how physical trajectories must occur. Meanwhile, our work establishes that awake generalized replays under TCM-SR dynamics can reverse directions and jump in physically unrealistic ways. Nevertheless, we can reconcile generalized replays with online planning through the theory developed in (Zhou et al., 2025). The TCM-SR theory identifies the TCM encoding update with the temporal difference learning update to the SR (Gershman et al., 2012). Sequential memory retrievals are equivalent to sampling from a weighted mixture of the SRs of the states represented in the current temporal context. The standard step-by-step rollout (from model-based reinforcement learning) is a special case when assuming Markov transitions and (*γ* = 0, *ω* = 0). In the general case of (0 < *γ* < 1), the sequence of memory retrievals is termed a “generalized rollout” since samples may jump ahead. We propose generalized replay as a neural correlate of generalized rollouts. In this framework, summing the values across a sequence of sampled (replayed) states provides an estimate of the initial state’s value for online evaluation. For intuition, the probability of sampling a state from the normalized SR is the correct expected discounted occupancy of a state in a standard rollout since rollouts simply sample transitions iteratively. Generalized replay is also reversible. In most natural environments, navigational state transitions are reversible. Even if only a forward transition is directly experienced, generalized replay can evaluate trajectories involving the reversed transition. Thus, TCM-SR theory explains how generalized replays can support efficient online evaluation of choices in the reversible sequential settings (Zhou et al., 2025).

Given the similarity between behavior time-scale free recall dynamics and replay timescale state transition dynamics, we propose a shared network mechanism that may underlie both. The bidirectional, forward-biased associations in human sequence memory has been connected to bidirectional hippocampal replay in rodents (Diba and Buzsáki, 2007; Zhou et al., 2026). We go beyond these works with detailed analyses of within-replay transition structure.

Previous models and experiments have placed temporal context in the rodent lateral entorhinal cortex (Howard et al., 2005; Tsao et al., 2018; Kanter et al., 2025) and human medial temporal lobe (hippocampus and amygdala) (Folkerts et al., 2018). We hypothesize that the hippocampal output region CA1 samples the next memory through a competitive process since CA1 predictive representations are consistent with encoding the SR (Stachenfeld et al., 2017), which was identified with the context-to-stimulus sampling matrix in the TCM (Gershman et al., 2012). The sampled state can influence the next retrieval by updating the temporal context in entorhinal cortex and CA3. The TCM-SR sampling account of generalized replays presents an alternative hypothesis for generating spatial trajectories, which was previously modeled as movement on the manifold of a continuous attractor network in the recurrent CA3 region due to firing rate adaptation (Azizi et al., 2013; Ecker et al., 2022; Mallory et al., 2025; Ji et al., 2026) or a joint entorhinal-hippocampal system (McNamee et al., 2021). Our account differs from the CA3-only models in that CA1 plays an important role in sampling the next states and predicts replay structure disturbance if CA1 is silenced. The model of (McNamee et al., 2021) shares much of the proposed circuitry, except with an emphasis on medial entorhinal cortex rather than lateral entorhinal cortex. However, the generation mechanism is a spectrally weighted diffusion propagator through continuous space, which differs from retrieving discrete states (as in jump and hover dynamics (Pfeiffer and Foster, 2015; Berners-Lee et al., 2022)) and does not predict asymmetric transitions, which we find in the W-maze data.

TCM-SR makes additional predictions for generalized replay by its link to episodic memory. First, it is a learning model that updates the SR association matrices incrementally. This implies generalized replays evolve with experience (Berners-Lee et al., 2022). A dataset with sufficient novel environment data could support a comparison between learning and non-learning versions of the TCM-SR transition model. Second, TCM-SR is a theory of replay as memory retrieval, so it makes predictions for replay transition dynamics in other non-spatial state spaces. In rodents, odor sequence tasks present an opportunity to study replay of non-spatial states (Manns et al., 2007; Shahbaba et al., 2022). Overall, our study illustrates that comparison of episodic memory and replay can yield rich insights into hippocampal function.

There are several limitations to our study. When comparing transition dynamics, an alternative explanation for TCM-SR winning model comparison is that replay events could be comprised of a mixture of continuous and fragmented modes (Denovellis et al., 2021; Zheng et al., 2025), and only the mixture of modes is described better by TCM-SR on average. We find this alternative unlikely since TCM-SR wins model comparison on events identified by spatial correlation and same-map activation criteria that implicitly filters out events with fragmented or stationary dynamics.

Another important limitation arises from the asymmetry results. The linear track dataset from (Pfeiffer and Foster, 2015) showed decoded jump distributions were reverse-biased for 2/3 rats (Figure S9**C**). Although we cannot make strong conclusions about the effect of task design, we speculate that the task demands of three arm delayed alternation in (Lustig et al., 2025) resulted in a greater proportion of forward replays. Nevertheless, the robust forward asymmetry observed in human free recall is only partially consistent with rodent hippocampal replay. Our study is focused on linearized state spaces because direction-selective hippocampal responses enabled us to decode directional transitions for the asymmetry component of the TCM-SR hypothesis. In linearized mazes, it is also easy to estimate the empirical movement transition matrix whereas open 2D arenas present the challenge of under-sampling state transitions. For 2D environments, TCM-SR-based decoding would be suitable if the animal is path-constrained such as in labyrinth, honeycomb, and other graph-structured mazes. Lastly, the present study focuses on awake replay in male rats. We did not include female rats due to the unavailability of relevant datasets to our knowledge.

Our work provides a principled framework to validate assumptions in replay decoding methodology. We demonstrate its utility by statistically justifying a generalized conception of replay and provide a theoretical interpretation for these events. Specifically, we characterize generalized replay with a transition kernel that allows for jumps and direction reversals, and we find these events are prevalent in a memory- and planning-demanding task. The dynamics model was derived from the TCM-SR theory of evaluation in sequential settings, leading us to propose generalized replay as a neural instantiation of generalized rollouts for online planning.

## 4. Methods

### 4.1. Temporal Context model and the successor representation

How does the brain use episodic memories to make decisions in a sequential regime where the outcome occurs after an extended sequence of events? TCM-SR provides a process model in the Markov decision process formalism of reinforcement learning. We represent T distinct items or states (spatial or otherwise) denoted s_1_, s_2_, . . . , s*_T_* as one-hot encoded vectors (**x**_1_, **x**_2_, . . . , **x***_T_*) in a state space *S*. The drifting temporal context integrates new items and exponentially decays the previous items with time. Letting t index the order of presented items, the context **c***_t_* is updated according to

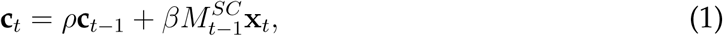

where *ρ*, *β* ∈ [0, 1] are parameters that set the drift rate and update strength, respectively. Adapting the model in (Sederberg et al., 2008), the context-to-stimulus matrix (initialized as a matrix of 0s) and stimulus-to-context matrix (initialized as the identity to represent pre-experimental context) are updated as:

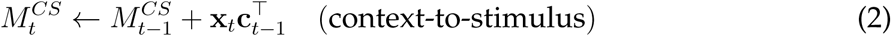

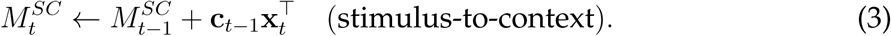

This process repeats until all items have been presented once. Explicitly, 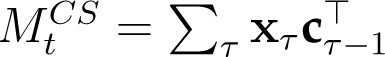 and 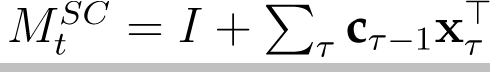.

At retrieval phase, the probability for retrieving the tth item is proportional to the similarity between the current retrieval context for the ith item **c***_i_* and its encoding time context **c***_t−_*_1_ (Figure S10**C**):

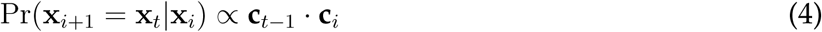

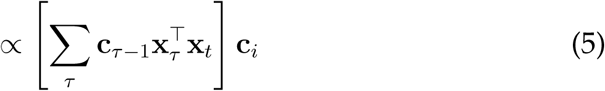

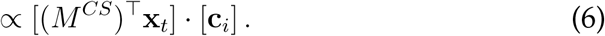

After each retrieval, the context is updated by eq. 1 (Figure S10**B**). The retrieval loop iterates until some termination condition occurs.

In (Gershman et al., 2012), the authors make a normative connection between TCM and the SR. For a given state space *S* and matrix of transition probabilities P(s*_t_*_+1|_ s*_t_*), the strictly forward-looking SR is defined as

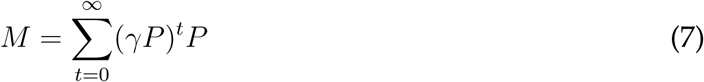

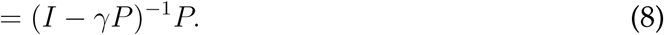

It is an infinite geometric series over future transitions to each state with a discount factor *γ* ∈ [0, 1) down-weighting the contribution of later transitions to the expected future occupancy. The additional factor of *P* makes the SR strictly future-looking.

Next, we re-derive the TCM-SR identification and derive an approximation used in this study. In (Gershman et al., 2012), the temporal context is shown to be equivalent to an eligibility trace for online TD(λ) learning of the SR. The future-looking SR update is (in row-start convention)

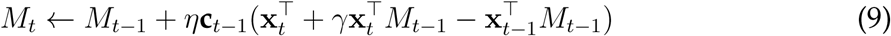

for eligibility trace **c***_t_* = *γ*λ**c***_t−_*_1_ + **x***_t_*. By comparing to eq. 1, the product *γ*λ was identified with the context decay rate ρ. If M *^SC^* = I, M = **0** at initialization and learning rate η = 1, then the first-pass update to each state is equivalent up to a transpose for the context-to-stimulus matrix and the SR, i.e. *M ^CS^* = *M ^⊤^*. This is because the last two terms satisfy 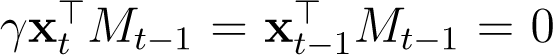 from the initialization. We refer the interested reader to (Gershman et al., 2012; Zhou et al., 2025) for further details and intuition.

Under this identification, the retrieval probability of the (i + 1)th state being the tth encoded state is:

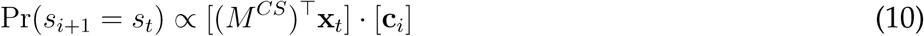

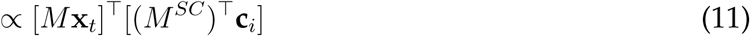

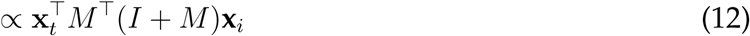

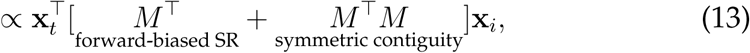

where we set (*M ^SC^*)*^⊤^***c***_i_* = (*I* + *M*)***x****_i_* if we assume *ρ_retrieval_* = 0*, β_retrieval_* = 1. This assumption essentially imposes Markov transitions in the context or hidden state. Finally, we can parameterize the relative strength of the forward biased SR component and the symmetric component by a weighting *ω ∈* [0, 1]:

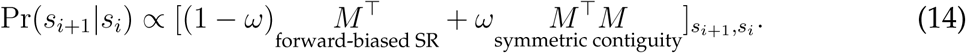

This retrieval probability distribution can be encoded as the transition matrix Pr(s*_i_*_+1|_ s*_i_*) in an HMM with parameters *γ* ∈ [0, 1), *ω* ∈ [0, 1]. See the second row of Figure 2**A** for how different choices of *ω*, *γ* affect the transition distribution. Note that this hidden state transition matrix is column-stochastic while most HMM software, including the ssm package we use, adopt a row-stochastic convention.

### 4.2. HMM transition models

We list alternative transition models in Table 2. The stationary, fragmented, and continuous kernels are used in (Denovellis et al., 2021). The movement kernel *P* is the empirical behavioral transitions during running at 100 ms time bins; we restricted the movement matrix to transitions within the same directional map to avoid ambiguity when the rat is at a track’s end. The strictly future looking SR was computed analytically using *M* = (I – *γP*)*^−^*^1^*P*. We include the reversed SR (Rev. SR) by transposing *M* and symmetrized TCM-SR (Sym. TCM-SR) by setting *ω* = 1 as a direction-reversed control and special case respectively since reverse replays are a known phenomenon. We also included an infinite *σ_c_* limit for the continuous kernel, which is uniform within the same map segment (Seg. Uniform).

**Table 2:**
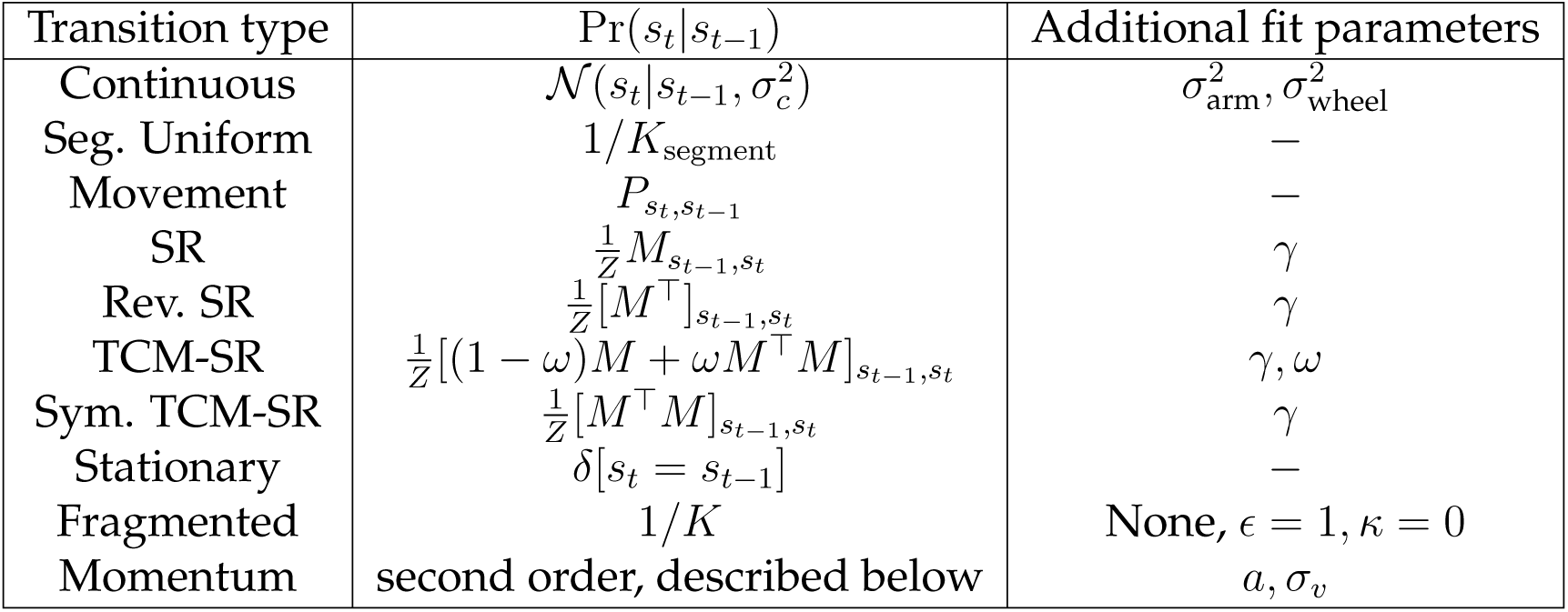
Transition matrices encoding different replay dynamics. Where present, *Z* represents the appropriate normalization factor to form a valid transition matrix. The Gaussian distribution of continuous transitions is discretized by spatial bins in each maze.

| Transition type | Pr(s<sub>t</sub>\|s<sub>t−1</sub>) | Additional fit parameters |
|---|---|---|
| Continuous | 𝒩(s<sub>t</sub>\|s<sub>t−1</sub>, σ<sub>c</sub><sup>2</sup>) | σ<sup>2</sup><sub>arm</sub>, σ<sup>2</sup><sub>wheel</sub> |
| Seg. Uniform | 1/K<sub>segment</sub> | – |
| Movement | P<sub>s<sub>t</sub>,s<sub>t−1</sub></sub> | – |
| SR | $\frac{1}{Z}M_{s_{t-1},s_t}$ | γ |
| Rev. SR | $\frac{1}{Z}[M^\top]_{s_{t-1},s_t}$ | γ |
| TCM-SR | $\frac{1}{Z}[(1-\omega)M+\omega M^\top M]_{s_{t-1},s_t}$ | γ, ω |
| Sym. TCM-SR | $\frac{1}{Z}[M^\top M]_{s_{t-1},s_t}$ | γ |
| Stationary | δ[s<sub>t</sub> = s<sub>t−1</sub>] | – |
| Fragmented | 1/K | None, ϵ = 1, κ = 0 |
| Momentum | second order, described below | a, σ<sub>v</sub> |

Beyond the kernel-specific parameters, we include additional fitted parameters for the first seven kernels: *κ* ∈ [0, 1] as the probability of self-transitions and *κ* ∈ [0, 0.02] as the residual probability of uniform transitions (0.02 is the probability of transitioning to the fragmented dynamic in (Denovellis et al., 2021)). We include a separate stickiness parameter κ because replayed locations have been shown to dwell between jumps, so separately parameterizing the probability of self-transitions can avoid distorting the kernel (Pfeiffer and Foster, 2015; Berners-Lee et al., 2022). We include ɛ to allow transient representations of other maps, which are otherwise forbidden by the transition matrices (Denovellis et al., 2021; Gu et al., 2026). Thus, for a given kernel type **T** ∈{continuous, movement, SR, . . .} , the actual transition matrix used in the sticky HMM is

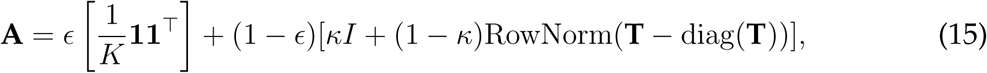

where **11***^⊤^* is the matrix of ones and **T –** diag(**T**) is **T** with diagonal entries set to 0. Lastly, we assumed a uniform initial state distribution in each event.

We have a separate transition dynamic for diffusion with momentum because it is second order Markov and infeasible to optimize parameters through EM. The transition probability is given by:

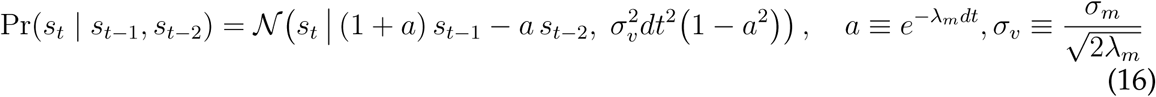

with parameters λ*_m_*, 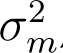, which specify a, σ*_v_*. The Gaussian kernel is discretized at bin centers. At the start of each trajectory, there is no s*_t−_*_2_ state, so we assume 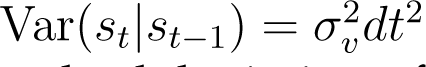 to avoid an additional parameter. σ*_v_* has units cm s*^−^*^1^ and scales the standard deviation of displacements (i.e. replay speed). *a* ∈ (0, 1) sets the correlation time scale, with 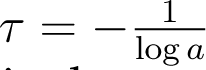 time bins. As *a* → 0, the kernel becomes memoryless, diffusive dynamics (equivalent to the continuous kernel). As *a* → 1, the kernel becomes a deterministic, constant velocity dynamic.

### 4.3. Datasets

We re-analyzed three prior extracellular recording datasets from dorsal hippocampal area CA1 of freely behaving rats on linearizable maze environments (Lustig et al., 2025; Grosmark and Buzsáki, 2016; Pfeiffer and Foster, 2015). All procedures in the original studies were approved by the respective institutional animal care and use committees (Janelia Research Campus, New York University, and Johns Hopkins University) and followed US National Institutes of Health guidelines on animal welfare. All subjects were adult male Long–Evans rats. Key properties of the three datasets are summarized in Table 3. Complete experimental details are given in the original reports and relevant information is summarized here.

**Table 3:** Summary of the three re-analyzed datasets.

|  | Lustig et al. (2025) | Grosmark and Buzsáki (2016) | Pfeiffer and Foster (2015) |
|---|---|---|---|
| Subjects | 3 male Long–Evans<sup>*a*</sup> | 4 male Long–Evans | 3 male Long–Evans |
| Weight at surgery | 300 g to 400 g | 250 g to 350 g | 450 g to 550 g |
| Age at surgery |  |  | 10–20 weeks |
| Behavioral task | W-maze with running wheel | novel linear track,<br>pre/post sleep | 1.8 m linear track |
| Electrodes | 4 silicon probes<br>(64-site, 6–8 shank) | 2 silicon probes<br>(64-site, 6–8 shank) | 40 tetrodes<br>(microdrive array) |
| Amplifier | Amplipex KJE-1001 | Amplipex (256-ch) | n/a |
| Sampling rate | 20 kHz | 20 kHz | 32 556 Hz |
| Spike sorting | KlustaKwik/Klusters | as in Harris et al. (2000) | xclust2, manual |
| Position tracking | 30 Hz | 39.69 Hz | 60 Hz → 30 Hz |
| Units analyzed here | 123–585 | 48-120 | 124-209 |
| Cell types included | pyr. + int. | pyramidal only | pyramidal only |
| Availability | contact corresponding authors | CRCNS (Grosmark et al., 2016) | contact corresponding authors |

<sup>*a*</sup>Data from 4 additional rats trained on a similar task, not published in Lustig et al. (2025), are also included.

#### W-maze dataset (Lustig et al., 2025)

Three adult male Long–Evans rats (300 g to 400 g at the time of surgery) were housed individually with running wheels on a 12 h light cycle. Rats were trained on a W-maze consisting of three parallel arms extending from a central hub, with automated doors controlling arm access, a brake-constrained running wheel in the hub, and lick-detector-gated water reward at the end of each arm. Rats alternated arms in the order left-center-right-center-repeat with wheel running between each trial to receive reward; the task is described in full in (Lustig et al., 2025). Four additional adult male Long–Evans rats (300 g to 400 g at the time of surgery) were trained on the same W-maze but a different three arm delayed alternation task with the order of arm alternation being left-center-right-repeat. This experiment is not described in (Lustig et al., 2025), but we refer to these data alongside the data from the first three as “W-maze” recordings. Each rat performed one session per day, typically ∼ 90 trials over 60 min to 75 min. Four silicon probes (8-shank 64-site Buzsaki64 or 6-shank 64-site Buzsaki64sp, NeuroNexus) were implanted over dorsal hippocampus through elongated craniotomies centered at AP –2.9 mm, ML ±2.2 mm and AP –4.2 mm, ML ±3.0 mm, and lowered to 1.0 mm to 1.5 mm beneath the brain surface. Spikes were detected above a threshold of the mean +1.5 SD of the high-pass filtered signal, automatically clustered with KlustaKwik, and manually refined in Klusters.

#### Novel linear track dataset (Grosmark and Buzsáki, 2016)

Four adult male Long–Evans rats (250 g to 350 g at the time of surgery) were water-restricted to 90% of their initial body weight beginning one week after surgery, with one day of ad libitum access per week. Before novelty sessions, rats were pre-trained to for-age for water reward on a geometrically unrelated open-field “cheeseboard” maze and were acclimatized for at least one week in the familiar room in which all sleep recordings were performed. Rats were implanted bilaterally in dorsal hippocampus with two 8-shank (Buzsaki64) or two 6-shank (Buzsaki64sp) silicon probes, oriented parallel to the septo-temporal axis and advanced over several days until CA1 was reached, as identified by sharp-wave ripples and pyramidal-cell activity and confirmed histologically post hoc. Each rat also carried a three-axis accelerometer to index movement during sleep. Hippocampal, EMG and accelerometer signals were recorded continuously at 20 kHz on 256-channel Amplipex systems (16-bit, analog multiplexing). Recordings were in the familiar room and two novelty rooms. Unit identification and spike sorting were performed by the original authors as described in Harris et al. (2000).

#### Open-field and linear-track dataset (Pfeiffer and Foster, 2015)

Three adult male Long–Evans rats (10 weeks to 20 weeks, 450 g to 550 g) were housed on a standard 12 h light cycle, handled daily, and food-restricted to 85–90% of free-feeding weight; behavioral training and recording were performed in the afternoon to early evening. Rats foraged for liquid food reward by running back and forth for reward on a 1.8 m linear track with a reward well at each end. Neural activity was recorded from a micro-drive array of 40 gold-plated tetrodes implanted in dorsal CA1. Units were identified by manual clustering on spike-waveform peak amplitudes using custom software (xclust2, M. A. Wilson), and putative inhibitory units were excluded on the basis of spike width and mean firing rate. Position was tracked at 60 Hz from two distinctly colored head-mounted LEDs by an overhead video system.

### 4.4. Common preprocessing

To make the three datasets comparable we applied the following steps uniformly.

#### Position and speed

Tracked position was down-sampled to a common 30 Hz before analysis. Instantaneous speed was computed as the distance between successive position samples in 2D divided by the sampling interval, and epochs were classified as “running” or “not running” by a threshold of 5 cm s*^−^*^1^.

#### Unit inclusion

Unit inclusion was not identical across datasets because cell-type classification was performed differently by each set of original authors. From Lustig et al. (2025) we included all sorted units, comprising both putative pyramidal cells and interneurons. There were 123-585 isolated units per session. From Grosmark and Buzsáki (2016) we included 48-120 isolated putative pyramidal cells per session. From Pfeiffer and Foster (2015) we included 124-209 isolated putative pyramidal units per session. In all three datasets, we did not apply a minimum spatial-information threshold.

### 4.5. Spiking data analysis

We analyze putative hippocampal pyramidal neurons in (Grosmark and Buzsáki, 2016; Pfeiffer and Foster, 2015) and do not exclude putative interneurons in (Lustig et al., 2025) (although we check that model comparison results are consistent when putative interneurons are excluded), inheriting the methods from each study.

#### Computing place fields

Directional place maps were computed using linearized 3.5 cm bins along each arm of the W-maze in (Lustig et al., 2025), 4 cm bins along the linear track in (Pfeiffer and Foster, 2015), and 2 cm bins along the linear track in (Grosmark and Buzsáki, 2016). For each state (direction × position), occupancy time and spike counts were summed during times when the animal was running ≥ 5 cm s*^−^*^1^. Wheel states in the W-maze were defined as 125 ms time bins during the 8-second wheel run as in (Lustig et al., 2025). The resulting rate maps were smoothed with a Gaussian kernel with σ =5 cm (3.5 cm for (Lustig et al., 2025)). This resulted in K = 372 *−* 390 states for the sessions in (Lustig et al., 2025), K *∈ {*160, 200*}* states for the sessions in (Grosmark and Buzsáki, 2016), and K = 80 *−* 92 states for the sessions in (Pfeiffer and Foster, 2015).

#### PBE detection

In our study, we focused on population burst events, which are commonly used candidate events for hippocampal replay (Davidson et al., 2009; Pfeiffer and Foster, 2013; Lustig et al., 2025). We computed the population spike counts in non-overlapping 1 ms (0.8 ms for (Lustig et al., 2025)) bins and smoothed with a Gaussian window of 20 ms SD. A bin was eligible if an animal was non-running (and not on the wheel for (Lustig et al., 2025)) during the task epoch. The smoothed population firing rate was z-scored and contiguous segments with at least one bin above 3 SD were considered. Lastly, gaps of 5 ms between contiguous segments were merged, and the merged events were thresholded against a minimum PBE duration (100 ms).

#### Memoryless, uniform prior Bayesian decoder

Let the place fields computed during running in each session have firing rates λ*_i_*(*s*) for unit i and state *s*. Let the decode time bin width be Δt and the population vector of spike counts for the tth time bin be y⃗*^t^*. The log posterior probability of a state under the memoryless, uniform prior Bayesian decoder is given by

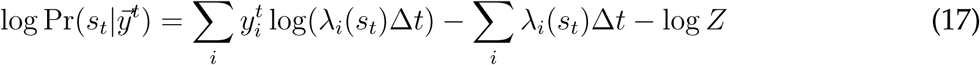

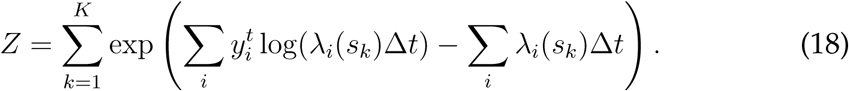

We clip the minimum of λ*_i_*(s)Δt to 10*^−^*^12^ for numerical stability, and low occupancy bins from the place map calculation are replaced with 0 in the log posterior space.

#### Traditional replays

We follow the weighted spatial correlation method for classifying replay events in (Lustig et al., 2025; Wu and Foster, 2014). In each PBE, we compute the MAP state over the entire concatenated state space (direction × position) in each decode window (20 ms bins stepping at 5 ms) under the memoryless, uniform prior Bayesian decoder. The time bin is excluded if less than 4 neurons fired in it or the peak posterior was less than 0.01. The surviving events in each of the 2 directional maps (10 maps in (Lustig et al., 2025) due to projecting 2D decoded locations to 3 arm polylines; see their methods) are scored by MAP-weighted Spearman correlation between time bin index and decoded position bin. Events were classified as “traditional replays” if the weighted Spearman correlation satisfied |ρ| ≥ 0.8 and had ≥ a8 steps in the same map. Once traditional replay time windows were computed, we rebinned the spikes into non-overlapping 10 ms time bins for the HMM comparison. We exclude events that occur on the wheel map in this event set.

#### Map-constrained trajectories

As a transition dynamic-agnostic alternative to spatial correlation criteria, we propose a threshold on contiguous time bins of the same map being active. During each PBE, we compute the posterior across the concatenated state space in 20 ms time bins stepping by 5 ms using the memoryless, uniform prior Bayesian decoder. We sum the posterior over each directional map and call the time bin “confident” if the total posterior probability on that map exceeds 0.6. In (Lustig et al., 2025), contiguous segments are merged over 5 ms gaps, and the merged segment is kept for downstream analysis if it exceeds 50 ms. In (Grosmark and Buzsáki, 2016; Pfeiffer and Foster, 2015), the single linear track only has two directional maps, so contiguous segments occur by random map activation more easily. Thus, we require 65 ms duration and do not merge across gaps. We rebinned the spikes into non-overlapping 10 ms time bins for the HMM comparison. We exclude events that occur on the wheel map in this event set.

### 4.6. Decoding error in synthetic data

We generate synthetic spiking data on the center arm of the W-maze using place fields derived from each session of the W-maze dataset under the continuous, momentum, TCM-SR, SR, and movement kernels. We restrict to the center arm to easily compute error distances on the same map of the maze. Spike counts are generated in non-overlapping 10 ms time bins with known ground truth hidden states, and the number of events and time bins are matched to the real data under the same-map activation criterion. Since the data was split into train (80%) and validation (20%) folds, we excluded 4 sessions with < 50 held-out time bins. The generative models used 100 random tuples uniformly sampled throughout the parameter space of each model. The stickiness parameter κ was each session’s fitted value in the real data and ɛ = 0 was fixed. Decoding models were fit on the training fold using either a parameter grid (momentum) or generalized expectation maximization (other fitted models) using the same procedure as on the real data (described in the next section). We compute the posterior using the forward-backward algorithm and calculate the RMSE between the posterior mean and the ground truth hidden state. This RMSE is compared to the RMSE of the oracle decoder, which refers to the original generative model at the true parameters. Finally, we compute the marginal log likelihood of each fitted decoder on the validation events and subtract from the oracle’s marginal log likelihood. This ΔNLL per bin is plotted against decoder error.

### 4.7. Transition model comparison

In all 3 datasets, we compared transition dynamics models on 3 different event sets: map-constrained trajectories at 10 ms/10 ms, all PBE bins at 3 ms/3 ms, and traditional replay events at 10 ms/10 ms (time bin/step). The map-constrained trajectory minimum duration was 50 ms in (Lustig et al., 2025) with 5 ms gaps merged and 65 ms in (Grosmark and Buzsáki, 2016; Pfeiffer and Foster, 2015) with no gaps allowed.

Sticky HMMs with different transition priors were parameterized as described in section 4.2. We performed cross-validated log likelihood comparison by randomly splitting events in each session into 5 folds. We fit parameters using the generalized expectation maximization algorithm (with exception of momentum dynamics, described in the next paragraph) for up to 200 iterations with exact E-steps and a gradient-based M-step (Dempster et al., 1977; Linderman et al., 2020). The M-step hyperparameters were: 25 iterations of gradient ascent on unconstrained parameters with learning rate 0.01 with each gradient clipped to (–0.05, 0.05). We added a Jeffreys prior Beta(0.5, 0.5) on the bounded parameters *γ*, *ω*, ɛ. We ensured convergence and enabled early stopping by tracking the change in marginal log likelihood per bin. Because we were comparing differences in transition dynamics, we did not recompute the place field observation model for different folds. The log likelihood on held-out folds was averaged for each session. To compute the fraction of events best described by each kernel, we computed the log likelihood on each event within a session using the models fitted on the full dataset.

The continuous transition hypothesis was encoded in a discretized Gaussian kernel with a capped standard deviation to preserve a notion of locality (Figure 3**A**, “Continuous”). The maximum standard deviations were 10.5 cm in the W-maze dataset and 12 cm in the linear track datasets, equivalent to 10.5 m s*^−^*^1^ and 12 m s*^−^*^1^ traversal speeds when decoding with 10 ms time bins. The 12 cm cap is more generous than the 7.3 cm standard deviation fixed in (Denovellis et al., 2021), so the best fit to spiking data could in principle support this choice.

Because the momentum dynamic is second-order Markov and only requires 2 parameters (a, σ*_v_*) or equivalently, (λ*_m_, σ_m_*), we grid searched the parameters for cross validated model comparison. We used log-spaced values

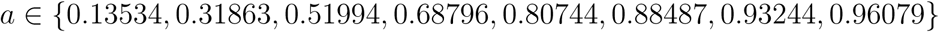

and

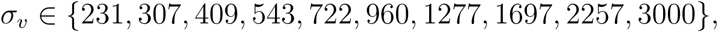

chose the highest log likelihood parameter combination on the train split, and evaluated on the validation split. Note that this range effectively capped *σ_v_* to 30 m s*^−^*^1^, but interior values of the grid were selected during model comparison. Lastly, we fixed *ɛ* = 0.01 and ignored κ in this model to keep the parameter grid tractable.

### 4.8. Model recovery

We conducted synthetic model recovery analysis on representative sessions from 3 rats in each dataset. First, we uniformly sampled 1000 points in parameter space (*ω*, κ, *γ*, σ_arm_*, σ*_wheel_*, ɛ*) for each transition model, dropping irrelevant parameters. Note that the stationary and fragmented kernels do not have parameters. Then, we generated synthetic spiking data using the place fields from each session for real event duration-matched time bins at 10 ms non-overlapping time bins. The same 5-fold cross validation model comparison procedure was used between the real and synthetic data, with the exception that the total EM iterations was capped at 50 instead of 200. For each synthetic dataset, the validation log likelihood was compared across models to call the recovered model class.

### 4.9. Jump distribution

The jump distribution was computed by taking the signed distance between decoded states during candidate events (MAP for memoryless, uniform prior Bayesian decoder, Viterbi path for HMM) within the same map. Hovers (0-bin changes) were excluded for visualization of the jumps. Jump signs were determined by the directional map the trajectory occurred on. The asymmetry index (“asym”) was computed as:

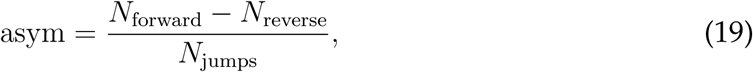

where N_forward_*_/_*_reverse_ was simply the number of displacements ≥+1 and ≤–1 bin, respectively. The p-value was a two-sided comparison against a transformed binomial null distribution Y defined by *X* ∼ Bin(*N*_jumps_, *p* = 0.5), *Y* = 2*X*/*N*_jumps_ –1.

Synthetic jump distributions were generated by uniformly randomly sampling an initial state and then sampling transitions using the fitted transition matrix **A** from the full data. The synthetic event lengths were matched to the event lengths in the real data. Again, hovers were excluded for visualization of the jumps.

### 4.10. Posterior predictive check on jump distribution

To evaluate the transition model prior, we compare these fitted priors from the full data to the one-step transitions in the neural data. We use memoryless Bayesian decoders with uniform prior to compute a jump distribution as described in the previous section using non-overlapping 10 ms time bins. Then, we generated synthetic trajectories in the hidden states as described above with initial states seeded at the real event’s first state, and we sampled spike-count matched observations. We apply the same memoryless Bayesian decoder with uniform prior to get *decoded* synthetic trajectories. We apply the same-map activation criteria to call events as used on neural data, and we resample synthetic trajectories of spikes until there are matched event counts to the real data, with 200 replicates. Jump distributions are computed from decoded synthetic trajectories. The percentiles (deciles marked) in each jump distribution (synthetic and from neural data) are plotted in the quantile-quantile plot.

### 4.11. Code and data availability

The paper analyzes data from (Lustig et al., 2025), (Grosmark and Buzsáki, 2016), (Pfeiffer and Foster, 2015). Data from (Grosmark and Buzsáki, 2016) is publicly available at CRCNS hc-11 (Grosmark et al., 2016). Please contact Brian Lustig and Albert Lee or Brad Pfeiffer and David Foster to obtain the respective analyzed datasets.

The code used to analyze the data and generate the figures will be released at https://github.com/george-tog/generalized_replay and the integration of TCM-SR into the hierarchical HMM package from (Denovellis et al., 2021) is available at https://github.com/george-tog/generalized_replay_classification.

### 4.12. Acknowledgments

We thank Brad Pfeiffer and David Foster for sharing their data with us and Andres Grosmark and György Buzsáki for making their data publicly available. This work was supported by the Kempner Institute for the Study of Natural and Artificial Intelligence (G.C., S.J.G.), the Department of Defense MURI program under ARO grant W911NF-2310277 (S.J.G.), a Polymath Award from Schmidt Sciences (S.J.G.), the Howard Hughes Medical Institute (B.L., A.K.L.), and the NSF CAREER award number 2543127 (M.G.M.). The authors have no competing interests to declare. We acknowledge the use of Claude Opus 5 and 5.5 to generate and refactor code in this study.

### 4.13. Author contributions

G.C., S.J.G., and M.G.M. conceptualized and designed the research. G.C. performed the analysis. B.L. and A.K.L. conducted additional experiments and preliminary analysis. G.C., S.J.G., and M.G.M. interpreted the results. G.C. wrote the initial draft of the manuscript. All authors revised the manuscript.

## Supplementary Information

**Table S1:** Cross-validation data budget for (Lustig et al., 2025) and additional four rats (decode bin 10 ms, step 10 ms, spatial bin 3.5 cm). Events are consecutive same-map activation based trajectory events. Train and validation columns are means over the 5 folds; Rat 2 rec01 has too few events for five folds, so it does not enter the main model comparison.

|  |  |  |  | Events per fold |  | Time bins per fold |  |
|---|---|---|---|---|---|---|---|
| Animal | Session | *N*<sub>folds</sub> | *N*<sub>events</sub> | Train | Validation | Train | Validation |
| Rat 1 | rec01 | 5 | 97 | 77.6 | 19.4 | 540.0 | 135.0 |
|  | rec02 | 5 | 105 | 84.0 | 21.0 | 532.0 | 133.0 |
|  | rec03 | 5 | 180 | 144.0 | 36.0 | 1001.6 | 250.4 |
|  | rec04 | 5 | 65 | 52.0 | 13.0 | 324.8 | 81.2 |
|  | rec05 | 5 | 99 | 79.2 | 19.8 | 539.2 | 134.8 |
| Rat 2 | rec01 | - | 4 | - | - | - | - |
|  | rec02 | 5 | 27 | 21.6 | 5.4 | 132.8 | 33.2 |
|  | rec03 | 5 | 14 | 11.2 | 2.8 | 66.4 | 16.6 |
|  | rec04 | 5 | 42 | 33.6 | 8.4 | 214.4 | 53.6 |
| Rat 3 | rec01 | 5 | 206 | 164.8 | 41.2 | 1228.8 | 307.2 |
|  | rec02 | 5 | 109 | 87.2 | 21.8 | 642.4 | 160.6 |
|  | rec03 | 5 | 131 | 104.8 | 26.2 | 818.4 | 204.6 |
|  | rec04 | 5 | 76 | 60.8 | 15.2 | 505.6 | 126.4 |
|  | rec05 | 5 | 136 | 108.8 | 27.2 | 791.2 | 197.8 |
|  | rec06 | 5 | 66 | 52.8 | 13.2 | 376.0 | 94.0 |
| Rat 4 | rec01 | 5 | 41 | 32.8 | 8.2 | 216.0 | 54.0 |
| Rat 5 | rec01 | 5 | 26 | 20.8 | 5.2 | 160.0 | 40.0 |
| Rat 6 | rec01 | 5 | 167 | 133.6 | 33.4 | 1004.0 | 251.0 |
| Rat 7 | rec01 | 5 | 24 | 19.2 | 4.8 | 181.6 | 45.4 |

**Table S2:**
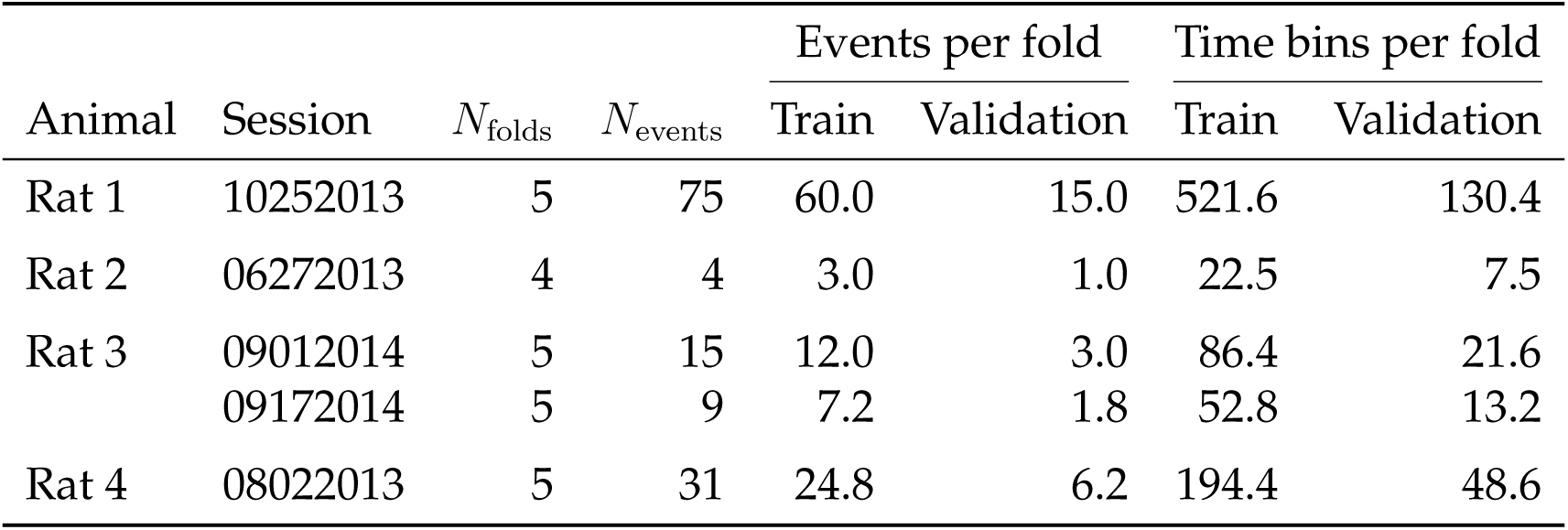
Cross-validation data budget for (Grosmark and Buzsáki, 2016) (decode bin 10 ms, step 10 ms, HMM bin 2 cm). Events are consecutive same-map activation based trajectory events. Train and validation columns are means over the *N*_folds_ folds; Rat 2 and Rat 3 session 09172014 had too few events for 5-fold CV on 4 parameters to be meaningful, so they do not enter the main model comparison.

|  |  |  |  | Events per fold |  | Time bins per fold |  |
|---|---|---|---|---|---|---|---|
| Animal | Session | *N*<sub>folds</sub> | *N*<sub>events</sub> | Train | Validation | Train | Validation |
| Rat 1 | 10252013 | 5 | 75 | 60.0 | 15.0 | 521.6 | 130.4 |
| Rat 2 | 06272013 | 4 | 4 | 3.0 | 1.0 | 22.5 | 7.5 |
| Rat 3 | 09012014 | 5 | 15 | 12.0 | 3.0 | 86.4 | 21.6 |
|  | 09172014 | 5 | 9 | 7.2 | 1.8 | 52.8 | 13.2 |
| Rat 4 | 08022013 | 5 | 31 | 24.8 | 6.2 | 194.4 | 48.6 |

**Table S3:** Cross-validation data budget for the (Pfeiffer and Foster, 2015) linear-track sessions (decode bin 10 ms, step 10 ms, HMM bin 4 cm). Events are consecutive same-map activation based trajectory events. Train and validation columns are means over the *N*_folds_ folds.

|  |  |  |  | Events per fold |  | Time bins per fold |  |
|---|---|---|---|---|---|---|---|
| Animal | Session | *N*<sub>folds</sub> | *N*<sub>events</sub> | Train | Validation | Train | Validation |
| Rat 1 | Linear1 | 5 | 124 | 99.2 | 24.8 | 956.8 | 239.2 |
|  | Linear2 | 5 | 215 | 172.0 | 43.0 | 1986.4 | 496.6 |
|  | Linear3 | 5 | 366 | 292.8 | 73.2 | 3606.4 | 901.6 |
|  | Linear4 | 5 | 78 | 62.4 | 15.6 | 596.0 | 149.0 |
|  | Linear5 | 5 | 139 | 111.2 | 27.8 | 1066.4 | 266.6 |
|  | Linear6 | 5 | 219 | 175.2 | 43.8 | 1863.2 | 465.8 |
| Rat 2 | Linear1 | 5 | 220 | 176.0 | 44.0 | 1952.8 | 488.2 |
|  | Linear2 | 5 | 242 | 193.6 | 48.4 | 2282.4 | 570.6 |
| Rat 3 | Linear1 | 5 | 203 | 162.4 | 40.6 | 2232.8 | 558.2 |

**Figure S1:**
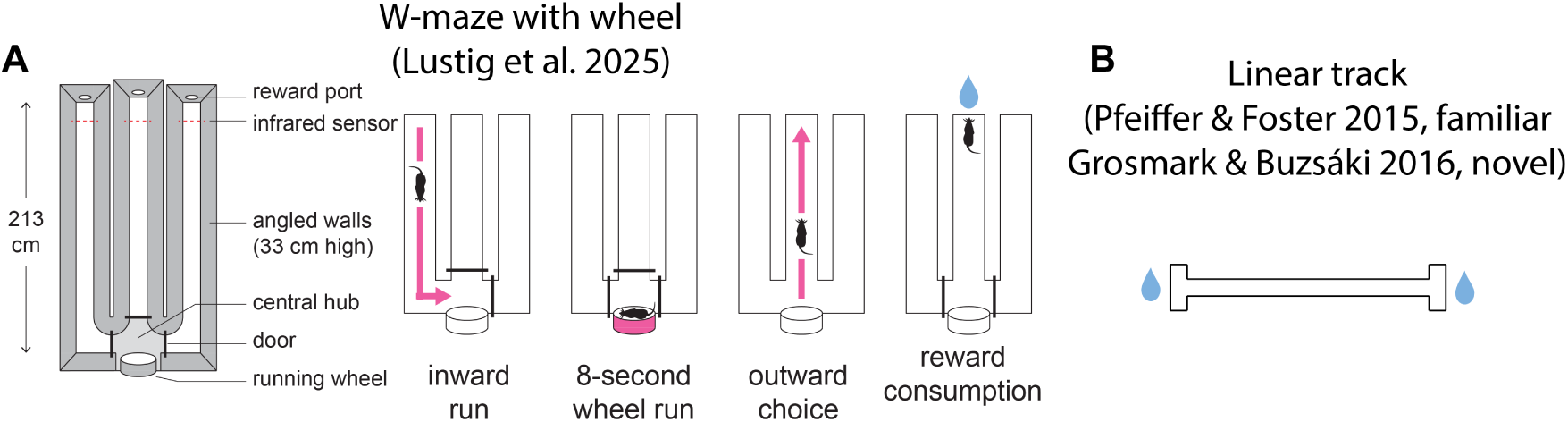
Task schematics for W-maze with delayed alternation and linear tracks. **A** Task schematic from (Lustig et al., 2025). Rats were trained to collect water reward at the end of each arm and alternate between arms in the following order: left→center→right→center→repeat (rats 1-3) or left→center→right→repeat (rats 4-7). Between each arm choice, the rat ran for 8 seconds on a wheel. **B** Task schematic of (Grosmark and Buzsáki, 2016; Pfeiffer and Foster, 2015). Rats were placed on a novel ((Grosmark and Buzsáki, 2016)) or familiar ((Pfeiffer and Foster, 2015)) linear track environment and collected water or liquid food reward at alternating ends of the track, respectively.

**Figure S2:**
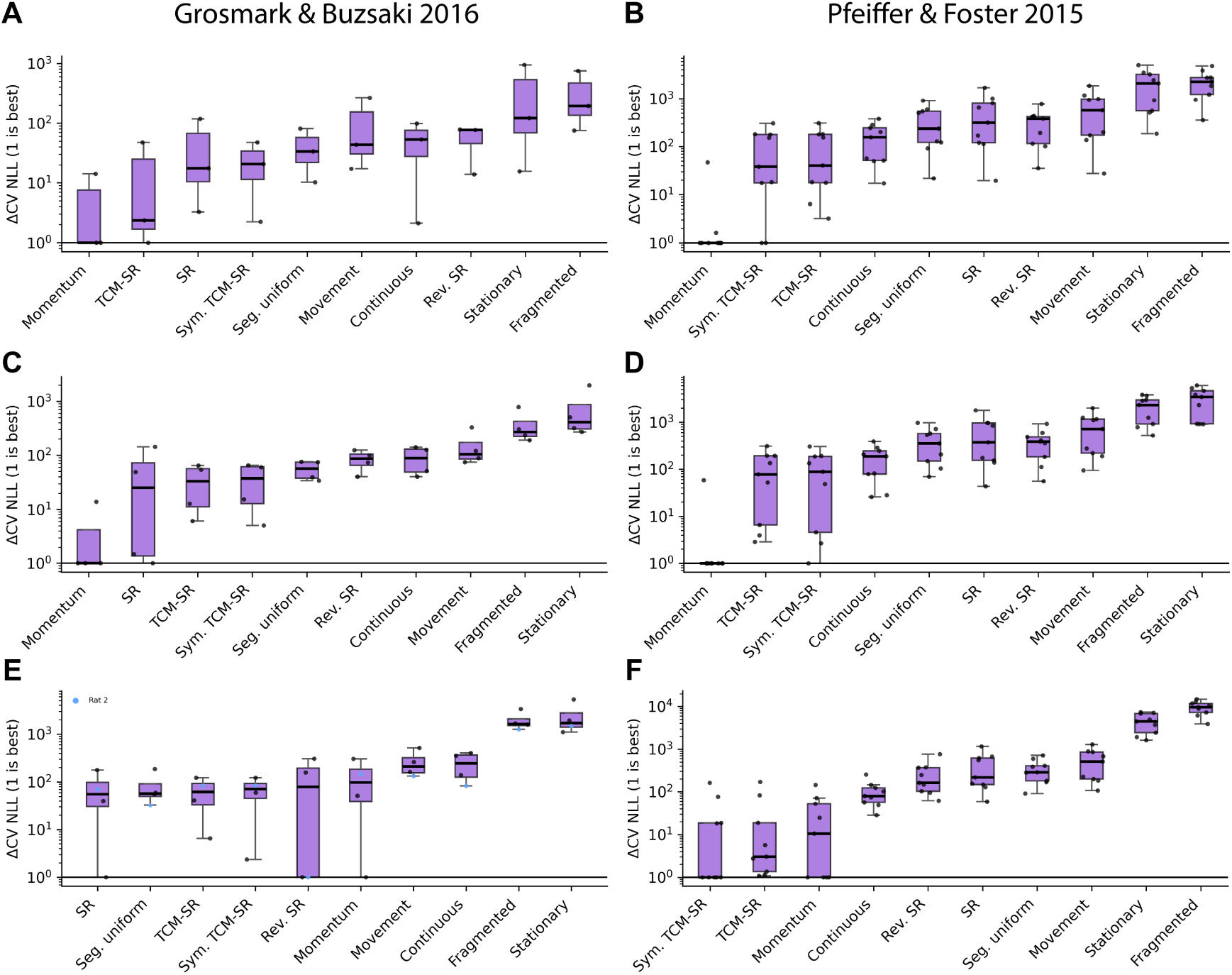
Replay dynamics on linear track are best explained by the diffusion with momentum transition model. In this figure, the left column corresponds to results from (Grosmark and Buzsáki, 2016) and the right column to (Pfeiffer and Foster, 2015). All panels depict difference in negative log-likelihood under 5-fold cross-validation at the event level. Each point denotes a single session. Events entering each comparison were **A-B** consecutive same-map activation at 10 ms time bins, **C-D** replay events identified by spatial correlations represented in 10 ms time bins, and **E-F** all PBEs in 3 ms time bins.

**Figure S3:**
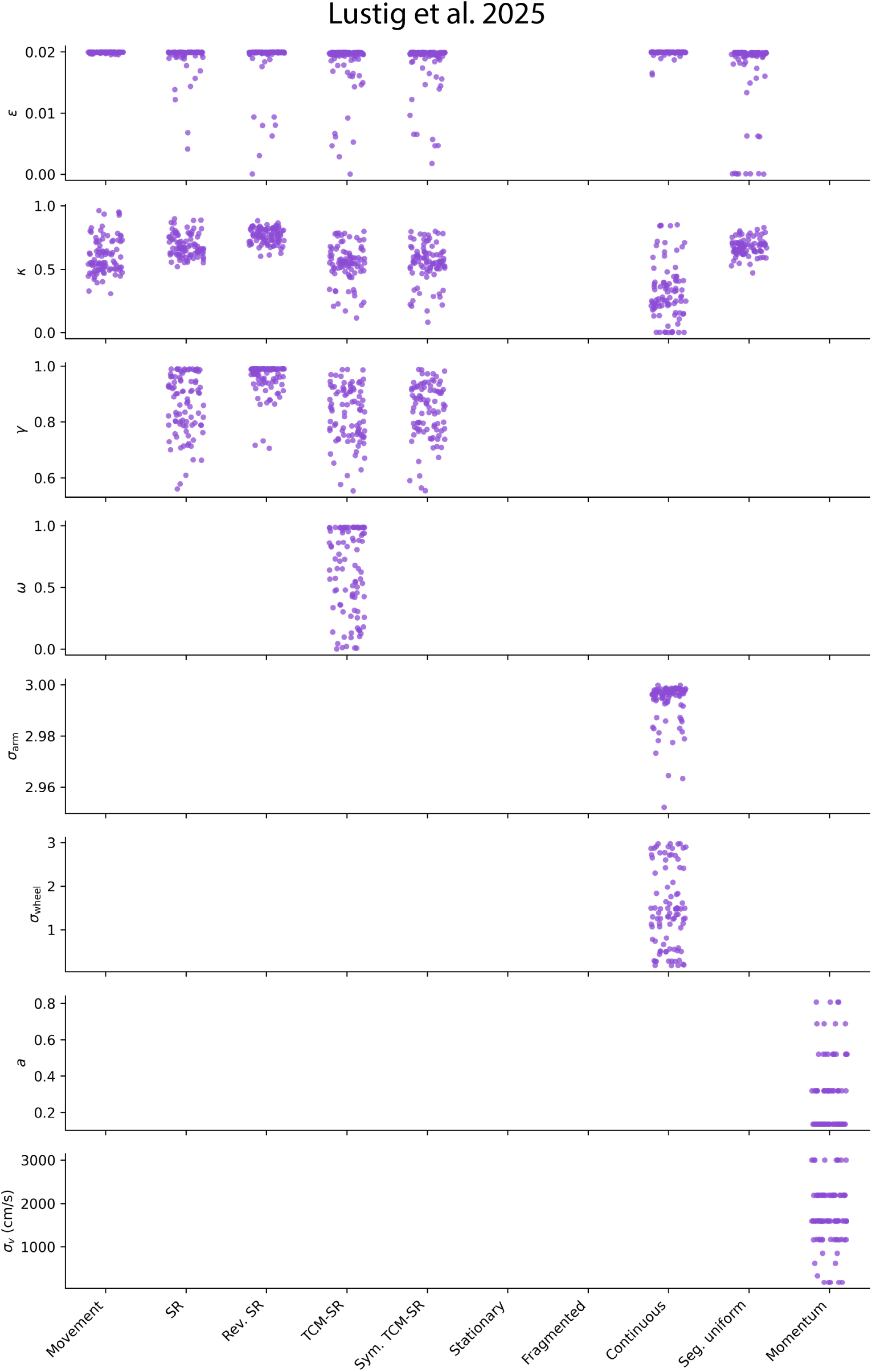
Fitted parameters in HMM transition kernels during 5-fold cross-validation. Each point represents the fitted parameter from one fold.

**Figure S4:**
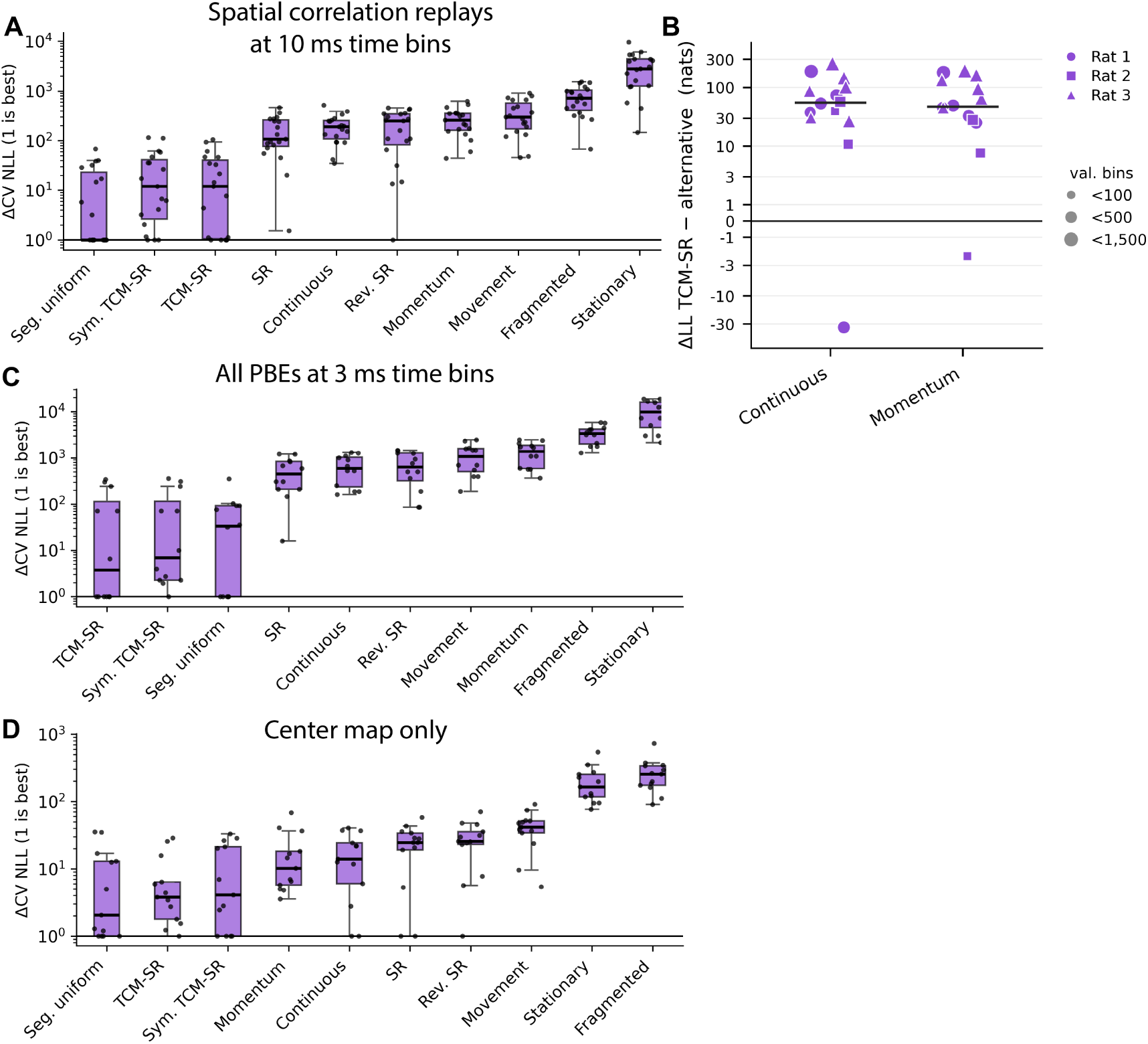
Transition model comparison is robust to event filtering, cross-session transfer, time bin size, and center arm subset. **A** Model comparison result on replay events identified by spatial correlation. **B** Average HMM log-likelihood after fitting on one session and evaluating on all other sessions for the same rat in (Lustig et al., 2025). The event set is same-map activation events and 10 ms time bins. There is one point per fitted kernel, and positive ΔLL favors TCM-SR. **C** Model comparison on all PBEs at 3 ms time bins. **D** Model comparison on same-map activation events subset to the center arm at 10 ms time bins.

**Figure S5:**
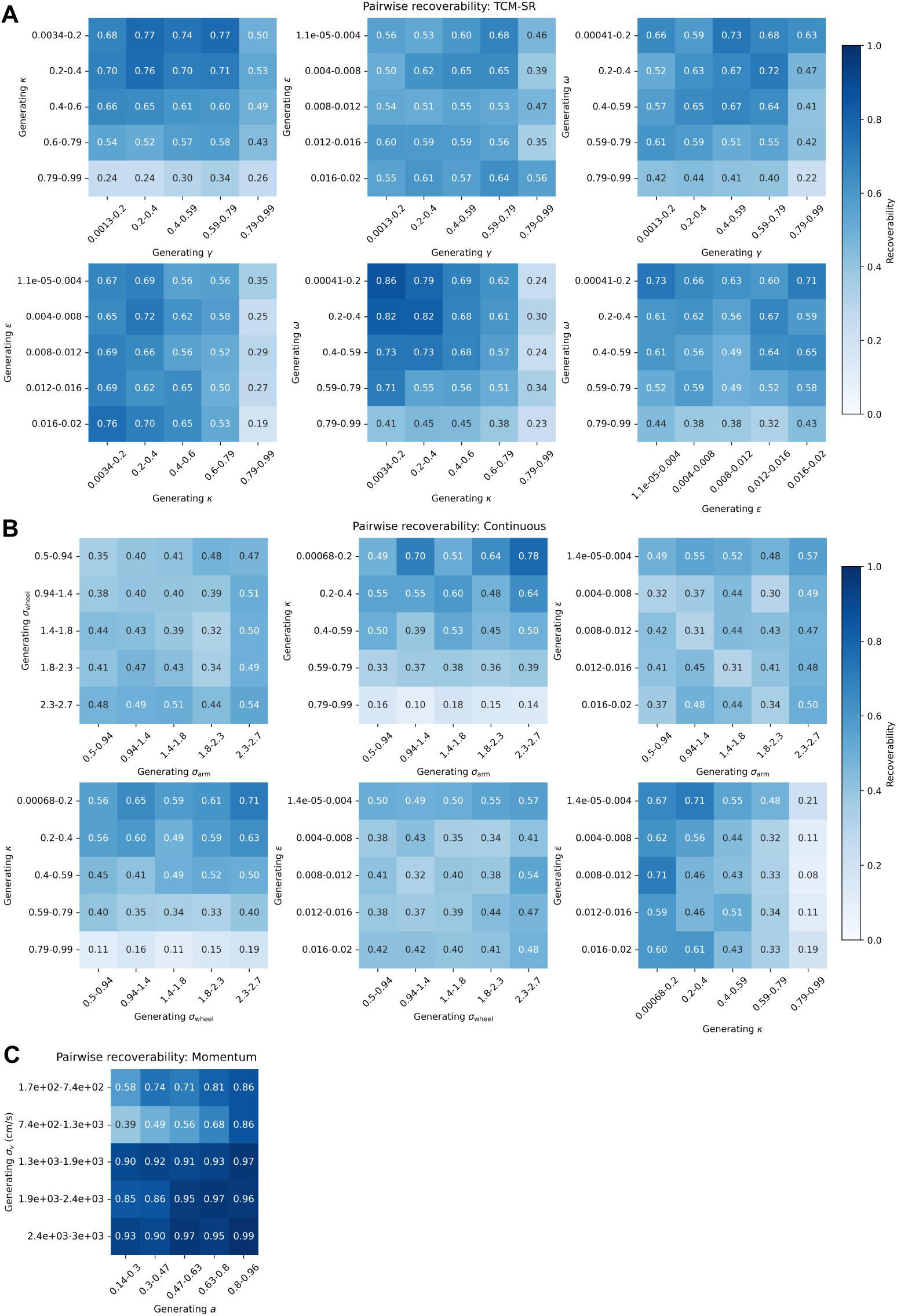
Synthetic recoverability of generating models across space of parameters. Recoverability for **A** TCM-SR, **B** continuous, **C** momentum kernel generating parameter value pairs.

**Figure S6:**
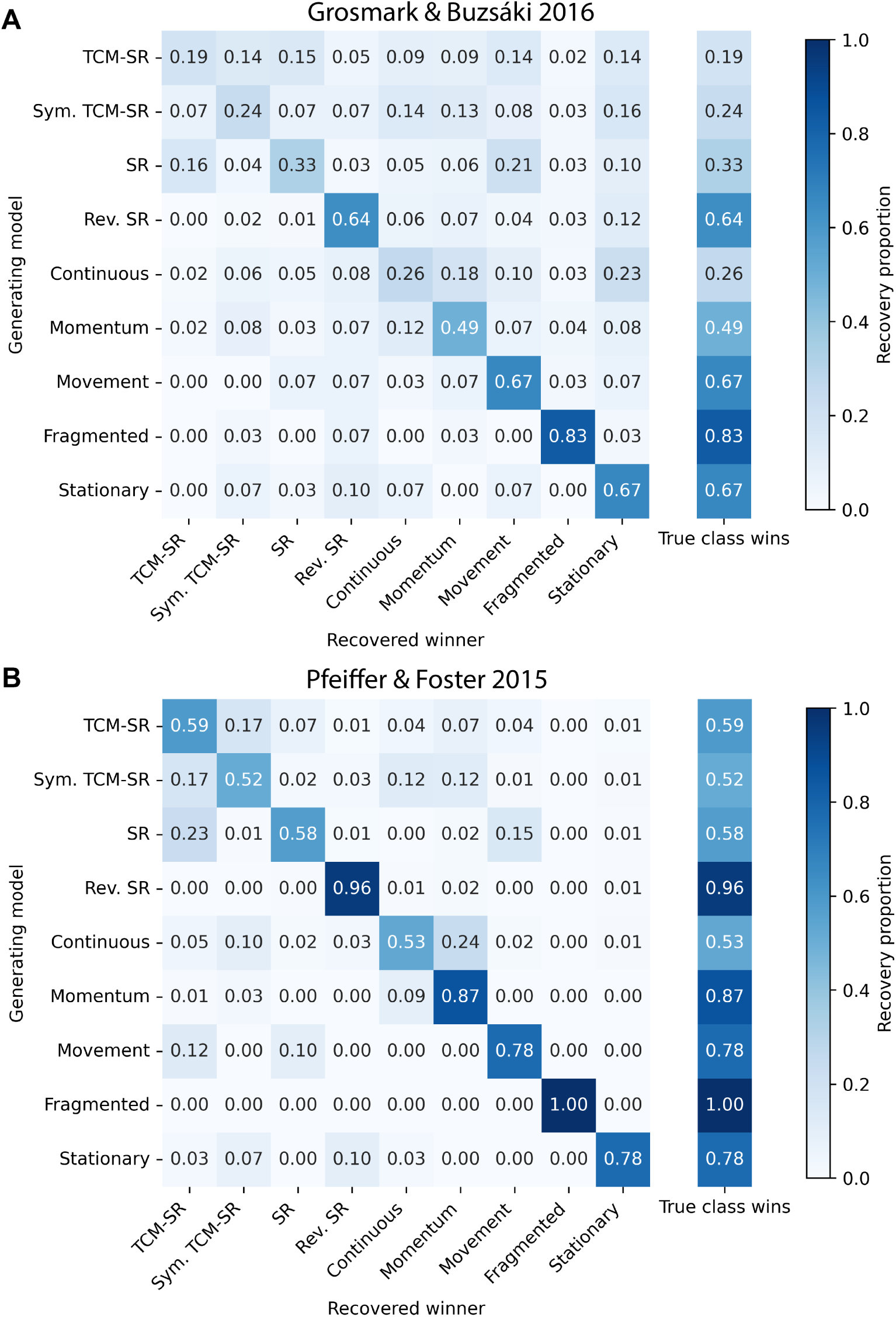
Synthetic recovery results for linear track datasets. Recovery matrix for **A** Grosmark and Buzsáki (2016) and **B** Pfeiffer and Foster (2015). For rats with multiple sessions, the session with the least event time bins was included in the recovery analysis.

**Figure S7:**
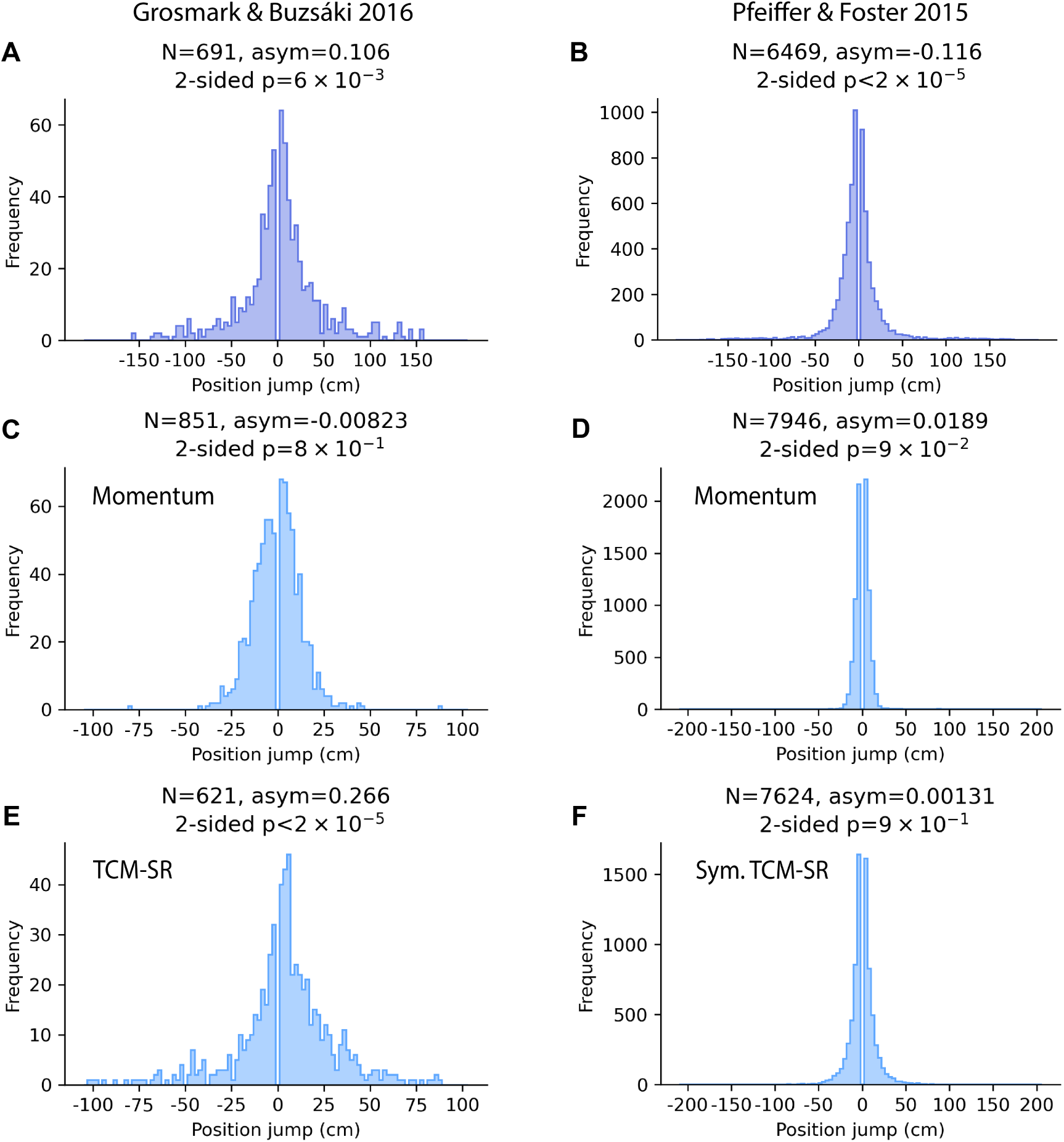
Jump distributions decoded from data compared to fitted priors for linear track data. Jump distributions under **A-B** memoryless Bayesian decoder with uniform prior from neural data (decoded at 20 ms non-overlapping time bins), **C-D** diffusion with momentum kernel sampled trajectories, **E-F** TCM-SR and symmetrized TCM-SR kernel sampled trajectories respectively.

**Figure S8:**
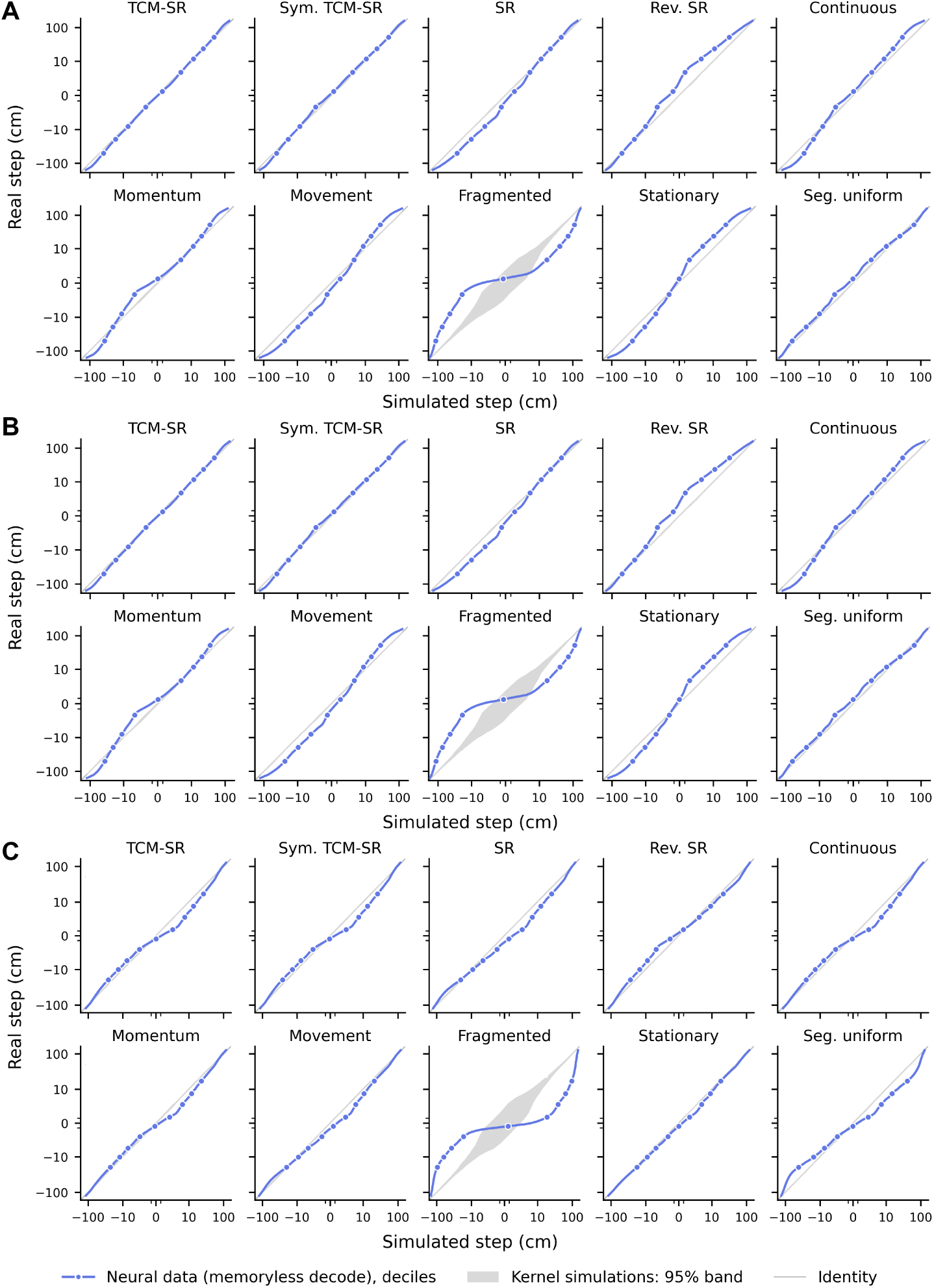
Quantile-quantile plots of jump distribution decoded from neural data and generated by fitted kernels. For each session, synthetic spikes were generated from each fitted transition kernel and filtered through the same-map activation criteria. Spike counts and time bins were matched to the real data. Then, using the same memoryless Bayesian decoder with uniform prior, MAP trajectories were decoded. Jump distributions are computed for the simulated data and neural recording. The blue line plots jump percentiles, points indicate deciles, and the gray band indicates the point-wise 95% band of the range of percentiles of synthetic jump distributions. The quantile-quantile plots are shown for **A** (Lustig et al., 2025), **B** (Grosmark and Buzsáki, 2016), and **C** (Pfeiffer and Foster, 2015). All models are different from the empirical jump distribution under Kolmogorov-Smirnov test at the 0.005 significance level.

**Figure S9:**
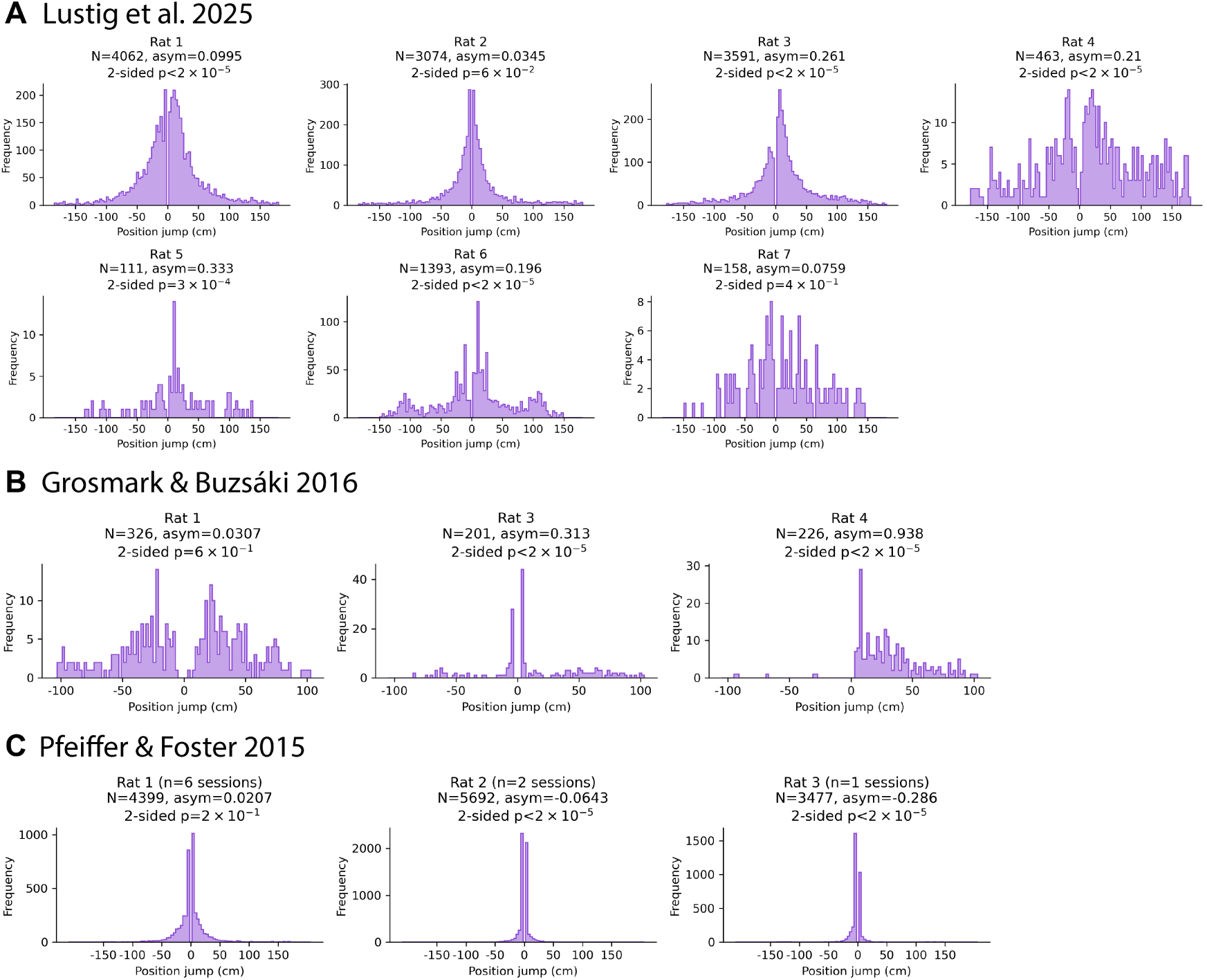
Jump distribution for each rat from Viterbi decoding with fitted TCM-SR or sym. TCM-SR kernel. Jumps were aggregated across all sessions for each rat in **A** (Lustig et al., 2025) and 4 additional left-center-right alternation animals, **B** (Grosmark and Buzsáki, 2016), and **C** (Pfeiffer and Foster, 2015) (decoded by sym. TCM-SR).

**Figure S10:**
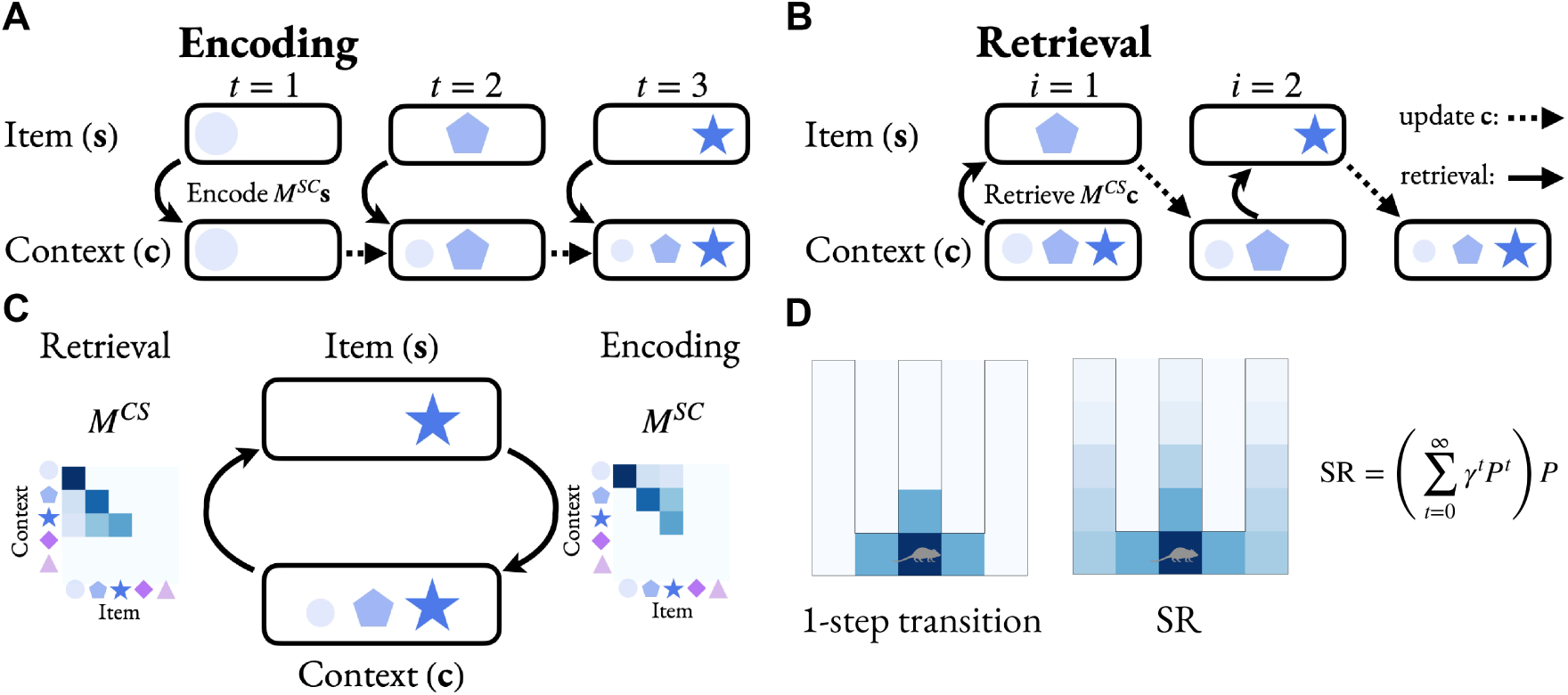
**A** TCM encodes items into memory in association with a drifting temporal context **c***_t_*. **B** TCM retrieves items based on the similarity of the current context to the context associated with each item at encoding time. The context is updated after each retrieval with the context associated with the just retrieved item. **C** As encoding proceeds, *M^SC^* and *M^CS^* are updated to encode item-to-context associations and context-to-item associations respectively. These matrices are necessary to retrieve items according to contextual similarity and update context upon retrieval. **D** Schematic of transitions and SR on a toy W-maze of grid states. *P* is a transition probability matrix between grid states under a policy that stays with some probability and chooses adjacent states uniformly. *γ* is a discount factor to down-weight states that are farther in the future. Left: one-step transition heatmap of transition matrix. Right: SR heatmap.

## Notes

### Competing Interest Statement

The authors have declared no competing interest.

